# Assembly of multi-subunit fusion proteins into the RNA-targeting type III-D CRISPR-Cas effector complex

**DOI:** 10.1101/2022.06.13.496011

**Authors:** Evan A. Schwartz, Jack P.K. Bravo, Luis A. Macias, Caitlyn L. McCafferty, Tyler L. Dangerfield, Jada N. Walker, Jennifer S. Brodbelt, Peter C. Fineran, Robert D. Fagerlund, David W. Taylor

## Abstract

CRISPR (Clustered regularly interspaced short palindromic repeats)-Cas (CRISPR-associated) systems are a type of adaptive immune response in bacteria and archaea that utilize crRNA (CRISPR RNA)-guided effector complexes to target complementary RNA or DNA for destruction. The prototypical type III-A and III-B CRISPR-Cas systems utilize multi-subunit effector complexes composed of individual proteins to cleave ssRNA targets at 6-nt intervals, as well as non-specifically degrading ssDNA and activating cyclic oligoadenylate (cOA) synthesis. Recent studies have shown that type III systems can contain subunit fusions yet maintain canonical type III RNA-targeting capabilities. To understand how a multi-subunit fusion effector functions, we determine structures of a variant type III-D effector and biochemically characterize how it cleaves RNA targets. These findings provide insights into how multi-subunit fusion proteins are tethered together and assemble into an active and programmable RNA endonuclease, how the effector utilizes a novel mechanism for target RNA seeding, and the structural basis for the evolution of type III effector complexes. Furthermore, our results provide a blueprint for fusing subunits in class 1 effectors for design of user-defined effector complexes with disparate activities.

**Important note:** While this manuscript was in preparation, a manuscript describing the structure of the type III-E effector was published^1^. We reference these important findings; however, a careful comparison of the structures will follow once the coordinates have been released by the PDB.

## Introduction

In the race for survival between bacteria and bacteriophage, CRISPR-Cas systems provide a bacterial adaptive immune response^2,3^. CRISPR-Cas effector complexes target foreign mobile genetic elements through sequence-specific hybridization with the crRNA guide and subsequent cleavage by Cas nucleases^4^. Type III CRISPR-Cas effectors target nascent RNA transcripts and cleave these with 6-nt cleavage periodicity^5,6^. Upon binding of an RNA target, type III systems initiate ssDNA cleavage using the HD domain of Cas10^7–9^. Furthermore, RNA target binding induces cyclic oligoadenylate (cOA) production by the palm domain of Cas10^10–13^. Cyclic oligoadenylates are allosteric activators of accessory nucleases, such as Csm6, which provide a second line of defense^10,11,14^.

Recently, two exciting studies characterized type III-E CRISPR-Cas systems^15,16^. The type III-E effector is composed of a single polypeptide made up of multiple Cas7 subunit domain fusions, including one domain split by a large insertion. Interestingly, this complex lacks Cas10 and Cas5, but still contains a Cas11 domain (Supplementary Fig. 1). These studies demonstrated programmable RNase activity by two of the four Cas7 domains and the architecture of this complex^13,17–19^.

A potential evolutionary intermediate between the multi-subunit type III-A/B systems and the single-subunit type III-E system is the type III-D system^15,20^. The III-D systems are marked by the presence of *csx10* (a specific variant of *cas5*), and often have a *csx19* gene, of which the function remains unknown. Previous reports have highlighted the evolutionary progression from multi-gene effectors (III-D1) to the single-subunit type III-E effectors^15^. Recently, a variant type III-D system (III-Dv) was described, showing multiple gene fusions, which suggest that it is positioned as an evolutionary intermediate between the multi-subunit and single-subunit effectors (Supplementary Fig. 1)^20^. Interestingly, the type III-Dv system has key differences compared to other subtype III-D systems, it maintains *csx19, cas10*, and *cas5* similar to III-D1, but includes a large insertion interrupting the terminal *cas7* gene, which appears conserved within type III-E systems^20^.

Despite a fascinating evolutionary relationship between these different type III effectors, there is a limited structural framework that illustrates this connection. Here, we present two cryo-electron microscopy (cryo-EM) structures of the type III-Dv surveillance complex (binary) and RNA target-bound (ternary) effector complex at 2.5- and 2.8-Å resolution, respectively. These structures shed important insight into the mechanisms of RNA targeting by these effectors. We demonstrate programmable RNA cleavage at three separate active sites across three unique Cas7 domains. Through analysis of structural rearrangements between the binary and RNA target-bound complex, we show how structural rearrangements allow for activation of the palm domain of Cas10 to prompt cOA production and ssDNA cleavage. By utilizing the recent power of Alphafold2, we predict a structural model of the type III-E effector complex from *Desulfonema isimotonii* (Cas7-11), which provides new insight into CRISPR-Cas evolution. Finally, the arrangement of subunits provided a scaffold for us to engineer a single-polypeptide III-D effector.

## Results

### The III-Dv effector forms a 332 kDa complex with no repeated subunits

The operon of the type III-Dv complex from *Synechocystis* contains *cas10*, a *cas7-cas5-cas11* fusion, a double *cas7* fusion (*cas7-2x*), *csx19*, and an insertion-containing *cas7* (*cas7-insertion*). Adjacent to the *cas* operon is *cas6-2a*, adaptation genes *and* a CRISPR array containing 56 spacers ranging in size of 34 to 46 bp (Fig. 1a)^21^. To determine the composition of the type III-Dv effector complex, we cloned the *cas* operon, *cas6-2a*, and first repeat-spacer-repeat of the CRISPR array from *Synechocystis* and expressed the operon in *E. coli*. The complex was purified using metal affinity and size-exclusion chromatography, where it eluted at ∼330 kDa (Supplementary Fig. 2). Analysis of the purified complex by SDS-PAGE and mass spectrometry confirmed the presence of all proteins except Cas6-2a, consistent with other type III complexes^5,22^ (Fig. 1c, Supplementary Table 1). The presence of Csx19 indicates this protein is a core component of the effector complex. Analysis of the crRNA length showed a mature crRNA of 37-nt, which agrees with a previous analysis of type III-Dv crRNAs in *Synechocystis* (Fig. 1b)^21^.

**Figure 1:**
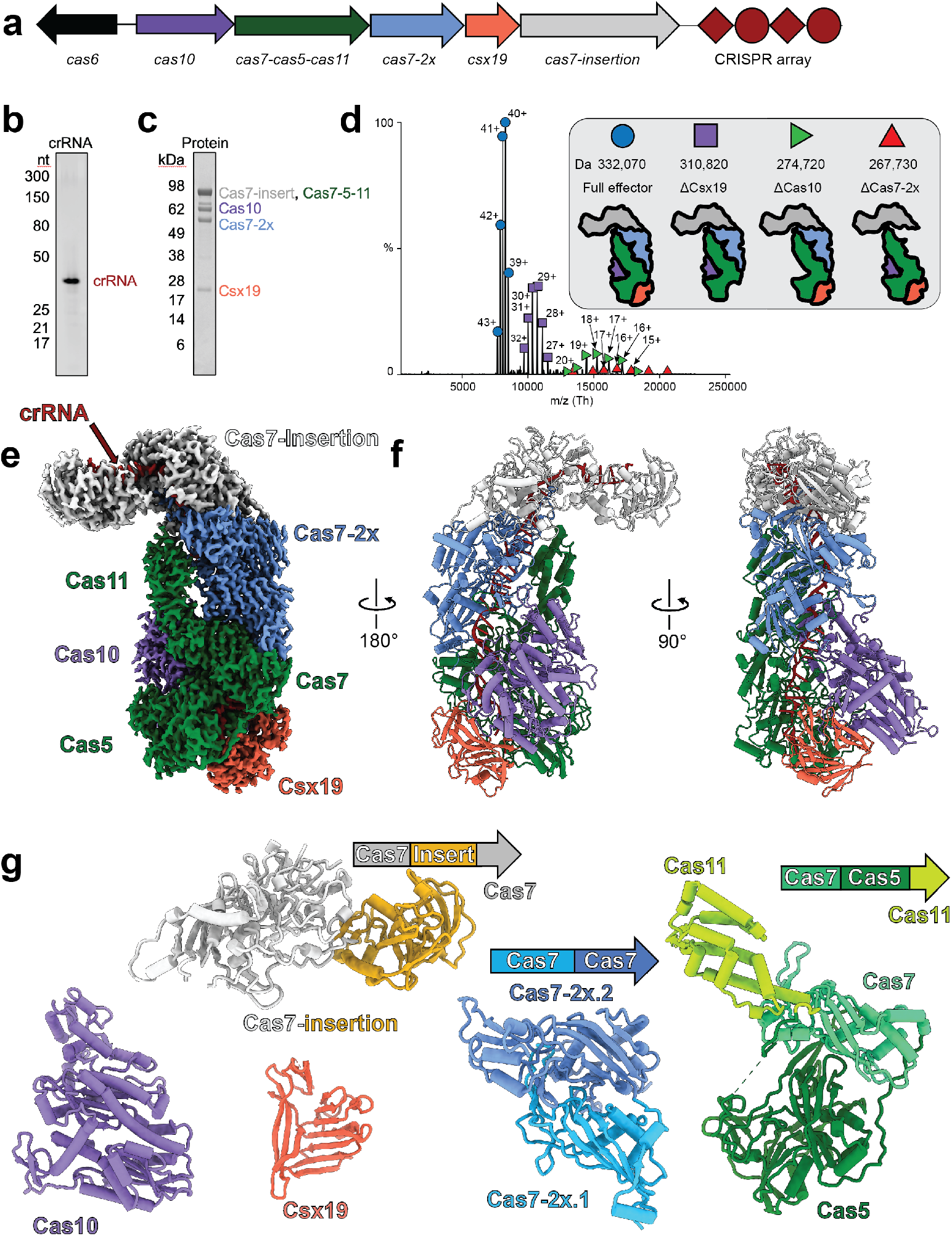
Stoichiometry and architecture of the *Synechocystis* CRISPR-Cas type III-D complex. **a**, Gene organization of the type III-D operon. Subunits and nucleic acids are colored as follows: *cas10*, purple; *cas7-cas5-cas11*, green; *cas7-2x*, blue; *cas7-insertion*, white. Cas6 is not found in the structure or purification and is thus colored black. **b**, TBE-Urea PAGE analysis of crRNA length for the type III-Dv complex. The crRNA product is 37-nt long. **c**, SDS-PAGE of the purified type III-Dv complex after size-exclusion chromatography. **d**, Native MS-MS of the type III-D complex. Peaks correspond to the full WT complex (circle), ΔCsx19 (square), ΔCas10 (green triangle), and ΔCas7-2x (red triangle). Ionization states are labelled for each peak. **e**, Cryo-EM map of type III-D binary complex. Subunits are colored as in the operon. **f**, Atomic model of the type III-D binary complex. **g**, Models of each subunit in the type III-D complex, colored by domains.

To confirm the composition and stoichiometry of this multi-subunit fusion protein effector, we performed electrospray ionization (ESI) native mass-spectrometry on the purified complex (Fig 1d)^23,24^. ESI showed one predominant peak corresponding to a native mass of ∼332 kDa, which is in excellent agreement with a complex composed of one Cas7-2x, one Cas7-ins, one Cas7-Cas5-Cas11, one Csx19, one Cas10, and a mature crRNA of 37-nt. Smaller peaks showed subcomplexes lacking either the Cas10, Cas7-2x, or Csx19 subunit, suggesting that these subunits are more likely to dissociate from the complex. The presence of each subunit was confirmed by subjecting the complex to denaturing top-down analysis with chromatographic separation (Supplementary Table 1). Biochemical and native mass-spectrometry analyses indicate that the type III-Dv effector can form subcomplexes. Based on this data, we hypothesize that Cas7-Cas5-Cas11 and Cas7-insertion assemble first, capping the two ends of the crRNA, followed by assembly of the rest of the subunits, Cas7-2x, Csx19, and Cas10.

### Structural analysis of the III-Dv binary complex

To delineate the architecture of this type III-Dv effector, we used cryo-EM to determine a 2.5-Å resolution structure of the complex containing the 37-nt crRNA (Fig. 1e, Supplementary Fig. 3e). We refined Alphafold 2-predicted structures of each subunit to rapidly generate a complete atomic model for the complex (Fig. 1f,g, Supplementary Fig. 4-6). After flexibly fitting and refining these subunits into the map, the overall architecture resembles a hammerhead shark, where the head is composed of the insertion containing Cas7, with the insertion domain and another small, uncharacterized domain creating each side of the head at the top of the complex (Supplementary Fig. 6). Interestingly, one side of the head (amino acids 1-112 of the Cas7-insertion N-terminus) was not observed in the cryo-EM map, likely due to flexibility. The body is composed of intertwined Cas7 and Cas11 domains of one Cas7-Cas5-Cas11 and one Cas7-2x subunit. Despite the Cas7 domains being part of larger fusion proteins or non-canonical subunits, they still arrange into a repeating backbone, wrapping around the crRNA and forming the major filament. This is a structural feature conserved across all class I effectors. At the base of the complex, Csx19 nestles next to Cas5, each forming one side of the tail. Cas10 forms the fin, sandwiched between Csx19 and Cas11, forming buried surface area with the bottom Cas7 and Cas5 domains. The placement of Cas10, Cas5, and Cas11 are typical of other type III CRISPR-Cas effector complexes.

This structure provides for a detailed understanding of how the domains of each of these fusion proteins are arranged. Interestingly, because the type III-Dv operon appears to have retained the domain organization of the type III-D1 operon, there are flexible linkers between the Cas7, Cas5, and Cas11 domains that allow for an unexpected structural organization of these subunits. A loop between the Cas7 and Cas5 domains (residues V221 to P244) and between the Cas5 and Cas11 domains (residues K602-T624) allows this fusion subunit to form an architecture that places Cas7 in the body of the complex, Cas5 below it at the tail, and Cas11 towards the head of the complex, on top of Cas7 – different to their arrangement in the operon (Fig. 1a,e). The purpose of utilizing long linkers between the domains may aid domain folding and complex assembly and may have been less evolutionarily expensive than re-arranging the genes to Cas5-Cas7-Cas11 before fusing them, which would allow for shorter linkers.

We utilized the Dali web server to search for structural homologues of each of the individual domains across the PDB^25^. Cas7 structural alignments revealed that all the III-Dv Cas7 domains aligned better with Csm3 (type III-A Cas7, average Z-score 16.7) than Cmr4 from (type III-B Cas7, average Z-score 12.3) (Supplementary Fig. 7a). However, structural alignments of III-Dv Cas5 domain (Csx10) revealed that it aligned slightly better with the Cas5 homologue of the III-B (Cmr3, Z-score 21.9, PDB 3X1L) than III-A complex (Csm4, Z-score 15.0, PDB 6xn7) (Supplementary Fig. 7b). Type III-Dv Cas10 also aligned better with III-B Cas10 (Cmr2, Z-score 21.4, PDB 3w2w) than III-A Cas10 (Csm1, Z-score 18.8, PDB 6o74) (Supplementary Fig. 7c). When aligned, the HD domain of the type III-A Csm1 hangs off the periphery of both the type III-Dv and III-B complexes (Supplementary Fig. 7d). Despite apparent loss of this HD domain based on Cas10 structural comparisons, bioinformatic work predicted an HD site for III-Dv in Cas10^20^. We were able to locate this putative site in our structure at residues H354, D355, and D356, indicating that type III-Dv Cas10 does indeed have an HD motif despite the absence of the canonical domain for this site present in Csm1 (Supplementary Fig. 7d). Our structure also maintains the conserved palm domain and GGDD motif of the Cas10 active site for cyclic oligoadenylate production from ATP. Running along the Cas7 major filament is the Cas11 domain of Cas7-Cas5-Cas11 and the C-terminus of Cas10, both of which are highly alpha-helical and resemble a small subunit. Interestingly, the Cas11 domain extends perpendicular to the trajectory of the Cas10 C-terminal domain, which is opposite to what is observed in type I and other type III complexes. Rather than a minor filament of repeating Cas11 subunits extending towards the top of the complex, type III-Dv utilizes two separate small subunit domains that extend from different domains into the minor filament. Together, these data clearly define the structural similarities of the III-Dv domains with known type III-A and III-B structures, despite many of these proteins being fused and the large insertion in the last Cas7 domain, as seen in the type III-E system^1^.

One notable difference between the type III-Dv complex and other multi-subunit type III complexes is the assembly of the Csx19 subunit. Csx19 is dominated by β sheets, and residue F71 caps the 8 nt 5’ crRNA handle through base stacking interactions between F71 and A1 of the crRNA (Supplementary Fig. 8a,c). R145 also contacts base G4 of the crRNA (−5 position in the 5’ crRNA handle) in a pocket containing the 5’-AAA-3’ tag of the crRNA (positions -2 to -4) (Supplementary Fig. 8d), suggesting a role of this subunit in stabilizing the 5’ end of the crRNA.

Despite these contacts, the role of Csx19 remains enigmatic. Interestingly, affinity purification of a ΔCsx19 complex with a N-terminal Cas10 tag did not result in the pulldown of complex, indicating that assembly of Csx19 onto the III-Dv complex is important for full complex assembly (Supplementary Fig. 9).

Another key difference is the presence of the Cas7-insertion subunit protruding from the effector complex, which has only been seen in the recent structure of the single polypeptide type III-E CRISPR-Cas effector^1^. In our structure, we found that the insertion domain within the Cas7-insertion subunit caps the 3’ end of the crRNA through base-stacking between F307 and A37 of the crRNA (Supplementary Fig. 8b). However, other than capping the 3’ end of the crRNA, the other roles of this domain remain unknown.

### Structural basis of RNA targeting by the III-Dv effector

To gain mechanistic insight into RNA targeting by this complex, we again utilized cryo-EM, solving the structure of the effector bound to target RNA at 2.8-Å resolution (Fig. 2a). The structure aligns nearly-perfectly to the binary structure. The RNA target hybridizes along the crRNA backbone using Watson-Crick base pairing and follows the same trajectory as the crRNA. Additionally, the two separate small subunit domains appear to stabilize the phosphodiester backbone of the RNA target using a positively charged surface, one from the C-terminus of Cas10 and one from the Cas11 domain of Cas7-Cas5-Cas11 (Supplementary Fig. 10). As in other class I complexes, every 6^th^ nucleotide of the crRNA and RNA target is flipped out by the β-hairpin thumb domain of each Cas7 domain, except for the Cas7-insertion subunit. Instead, this subunit threads an ordered loop through the crRNA-RNA target duplex. This loop pushes the stacking bases apart, between bases U29 and C30 of the crRNA and G27 and A28 of the RNA target. The lack of a protruding β-hairpin from the Cas7-insertion subunit causes a more rounded kink in the RNA target. This kinked position is only 4-nt upstream of the kinked RNA backbone of the closest Cas7 in Cas7-2x.2, a feature that has not been observed in multi-subunit type III complex structures.

**Figure 2:**
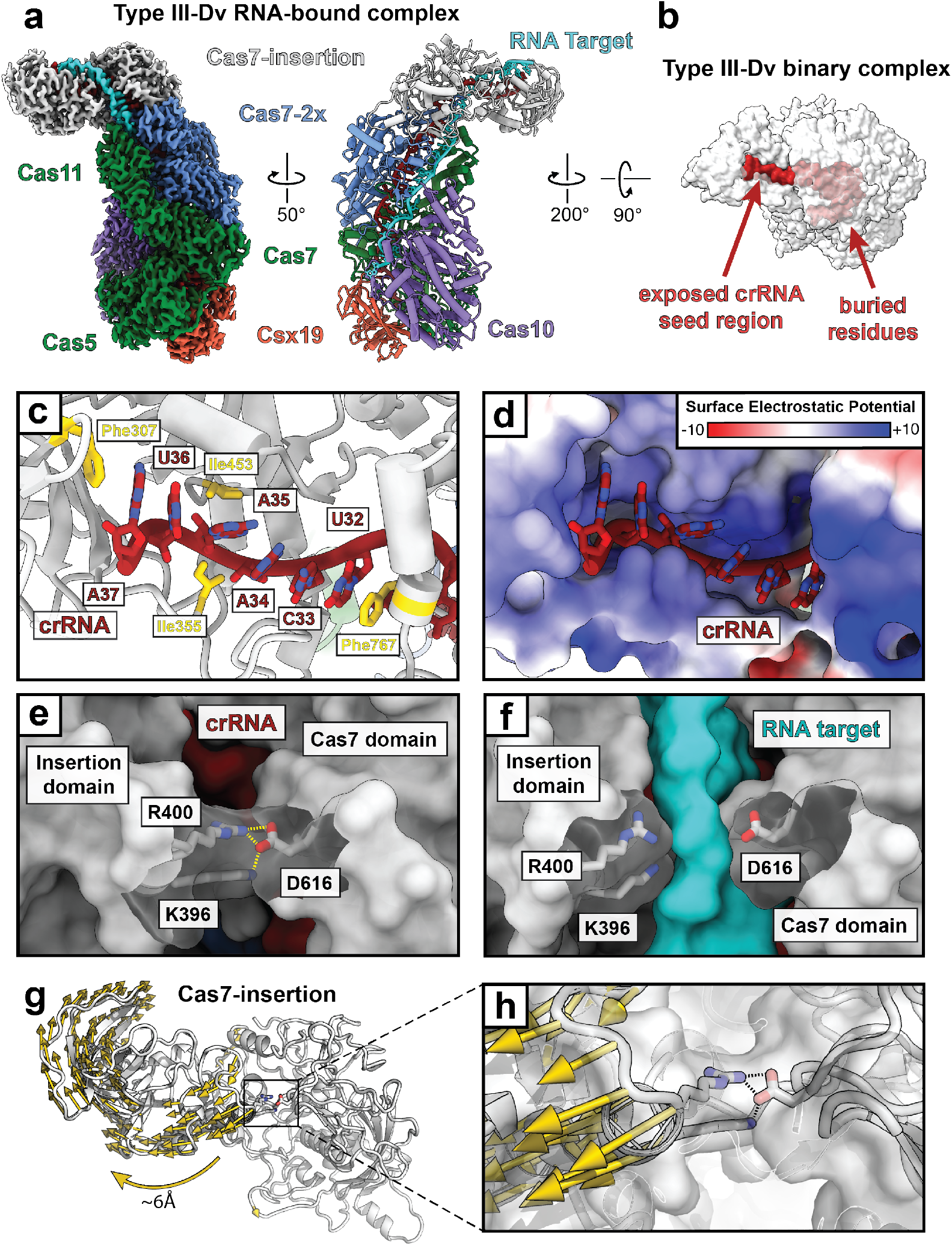
crRNA seed region initiates RNA target binding. **a**, Structure of the type III-Dv (ternary) complex bound to a target RNA, with the cryo-EM map on the left and atomic model on the right. **b**, Surface representation of the type III-Dv complex highlights the buried surface of the crRNA with the exception of a seed region that is exposed by the insertion domain of Cas7-insertion. Units are in kT/e. **c**, Exposed residues in the crRNA seed region stabilized by residues in the insertion domain. **d**, The crRNA seed region sits in a positively charged pocket of the insertion domain. **e**, A salt bridge between D616 and R400/K396 blocks RNA target binding at this region, presumably requiring seeding first. Dashed lines show the salt bridge interactions. **f**, Separation of the salt bridge in **e** to accommodate the RNA target. **g**, Modevector map showing the conformational change in the insertion domain of Cas7-insertion, allowing the salt bridge to break apart, as seen in **g**.

The type III-Dv binary structure reveals how the insertion domain of the Cas7-insertion subunit serves as an anchor that pulls the 3’ end of the crRNA spacer into a much different geometry than other type III and type I systems (Fig. 2b, Supplementary Fig. 8a,e). We observe the crRNA to be buried within the protein subunits throughout the entire complex with exception of a small region at the 3’ end of the crRNA that is anchored by the insertion domain (Fig. 2b). These six terminal bases of the crRNA (U32 – A37) lie in a positively-charged pocket of the insertion domain, positioning the Watson-Crick face of the bases towards the surface, primed for base pairing with an RNA target (Fig 2c,d). Base A37 is capped by F307 and base U32 forms a base-stacking interaction with F767, while I355 and I453 hold the seed region in place through hydrophobic contacts. A salt bridge between R400, K396, and D616 within the Cas7-insertion subunit joins the cleft between the insertion domain and the Cas7 domain, blocking RNA hybridization with the crRNA (Fig. 2e). Fascinatingly, this salt bridge breaks apart upon RNA target binding (Fig. 2f). We hypothesize that the six 3’ terminal bases of the crRNA serve as a seed region for initial binding of an RNA target. Sufficient hybridization between this crRNA seed region and an RNA target initiate conformational changes to open the salt bridge for continued RNA hybridization. Indeed, after analyzing the conformational changes within the Cas7-insertion subunit upon RNA binding, we see a significant shift in the insertion domain, swinging outwards to open the cleft between the insertion domain and Cas7 domain, breaking the salt bridge (Fig. 2g,h). Together, these results highlight a unique RNA target seeding mechanism among type III effectors. We predict that removal of this insertion domain removes the salt bridge shielding the crRNA from hybridization, likely abrogating the necessity of seed binding.

### Programmable target RNA cleavage by the type III-Dv effector complex

Next, we investigated the activity of the type III-Dv effector. Incubation of the complex with a 5’-fluorescently-labelled 60 nucleotide RNA substrate revealed cleavage of the RNA at positions 31, 37 and 43 nucleotides from the 5’ label (Fig. 3a-d). Interestingly, digestion of the same substrate but with a 3’ fluorescent label revealed only one predominant cleavage event positioned 17 nucleotides from the label (or 43 nucleotides from the 5’ end), suggesting a faster rate of cleavage at this position (Fig. 3c,e). Cleavage was metal-dependent, with optimal cleavage occurring with Mg^2+^ and Mn^2+^ (Supplementary Fig. 11). Interestingly, cleavage was observed almost immediately (Fig. 3d). These results reveal three active Cas7 domains that rapidly cleave target RNA and hint that they may differ in kinetics.

**Figure 3:**
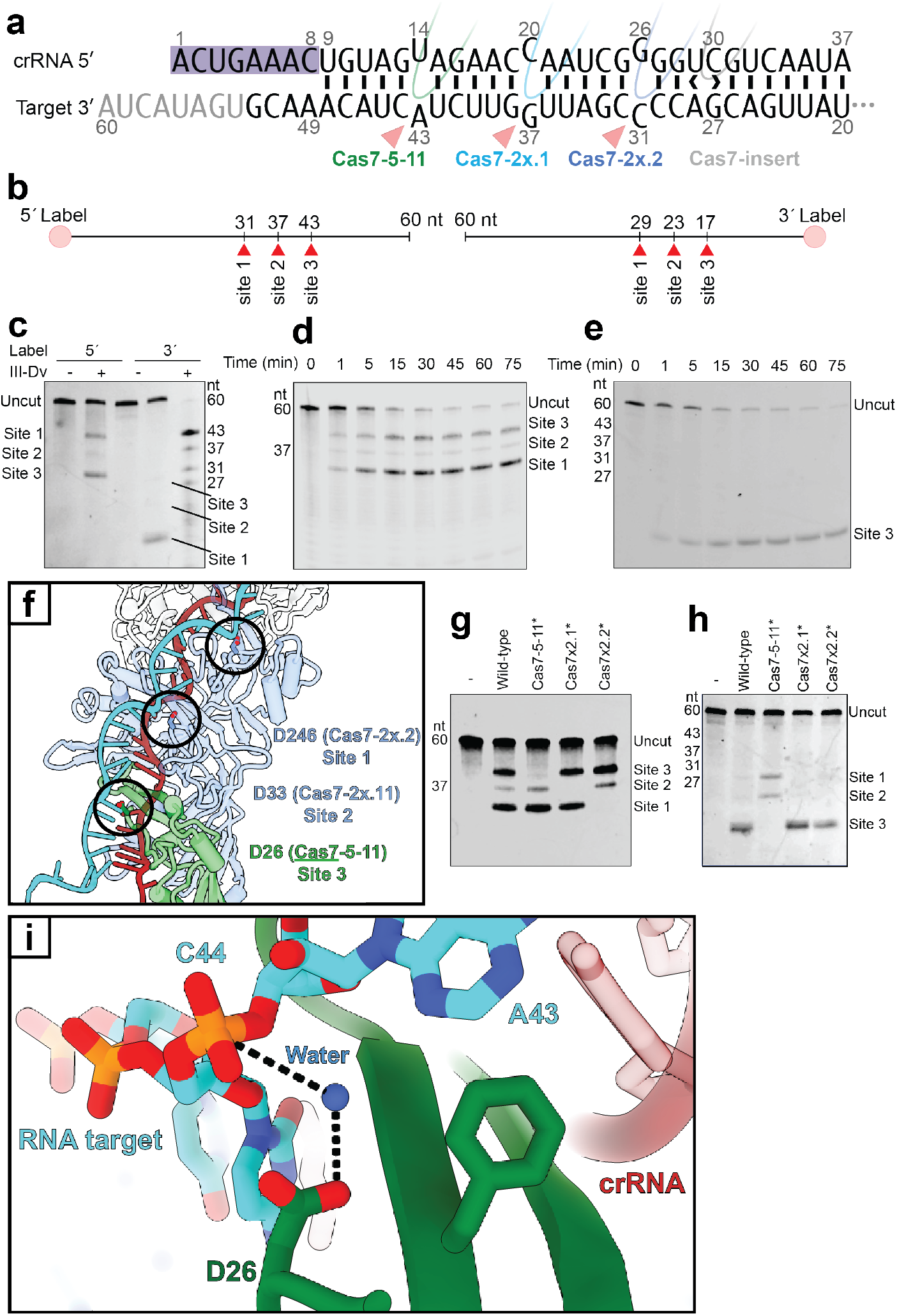
RNA targeting by type III-D. **a**, Representation of the crRNA-RNA target duplex. The observed bases (black) and 3’ bases of the target not tracible (gray) are shown. The 5’ bases of the target not tracible are not shown. The crRNA 5’ handle is highlighted (purple shade). Cleavage positions (red arrows) and respective catalytic Cas7 domains are indicated. **b**, Schematic of labeled RNA targets and cleavage locations. **c**, Cleavage of the RNA target with a 5’-FAM and 3’-FAM label. Products are visualized via TBE-Urea PAGE. **d**, RNA cleavage time course with a 5’-IRD800 labelled RNA target across 75 minutes. **e**, RNA cleavage time course with a 3’FAM labelled RNA target across 75 minutes. **f**, Active site aspartates for each Cas7-Cas5-Cas11 and Cas7-2x, each positioned at the kinked scissile phosphate. **g**, 5’IRD800-labelled and **h**, 3’FAM-labelled RNA cleavage analysis after mutagenesis of the three active site residues of Cas7-Cas5-Cas11, Cas7-2x.1, and Cas7-2x.2. **i**, Active site of Cas7-Cas5-Cas11 for ssRNA cleavage. Dashed arrows show the coordination of a putative water molecule.

To gain a structural and mechanistic understanding of RNA cleavage by this effector, we first scanned the structure for acidic residues positioned at the kinked phosphodiester backbone of the RNA target. Structural analysis of the Cas7 domains revealed aspartate residues positioned between the scissile phosphate of the target RNA and the β-hairpin thumb. These residues correspond to D26 of Cas7-Cas5-Cas11 (position 43 of the target), D33 of Cas7-2x.1 (position 37 of the target), and D246 of Cas7-2x.2 (position 31 of the target) (Fig. 3f). Interestingly, we found density at each active site that is positioned between the identified aspartate residue and the scissile phosphate (Supplementary Fig. 12a-c). Considering these densities were not present in the binary structure and remained unaccounted for after the full III-D-target complex was built, as well as the fact that divalent cations were not added to the buffer before freezing, we have putatively assigned these densities as water molecules. We hypothesize that these water molecules are coordinated by the given aspartate residue and primed for nucleophilic attack of the scissile phosphate of the target RNA. Despite no involvement from Cas7-insertion in RNA target cleavage, we observe density for a putative water molecule coordinated by an aspartate, positioned outside of the RNA backbone (Supplementary Fig. 12d). Further structural analysis with necessary metal ions is necessary to understand why this site does not participate in RNA cleavage.

To confirm the predicted active residues in the three active Cas7 domains, we mutated each aspartate to alanine, expressed and purified each variant, and tested these for cleavage activity against 5’ and 3’ fluorescently labelled RNA substrates (Fig. 3g,h). Mutation of the predicted active aspartate residues in the Cas7 domains successfully disrupted each cleavage event independently of the other. This could allow programming at these discrete and independent cleavage sites and exploited to create a III-Dv effector as a programmable sequence-specific RNase enzyme.

### Conformational changes activate Cas10 for cOA production

We next sought to understand whether the III-Dv complex retains a secondary immune response through activation of Cas10 upon non-self RNA target binding. In our structure, while the target RNA engages in Watson-Crick base pairing along almost the entirety of the crRNA, after position C8 in the crRNA, the target RNA disengages at the anti-tag sequence and is funneled into an exit channel on the surface of Cas10 (Fig. 4a). This is reminiscent of non-self-targeting that occurs within the Cas10 subunits of type III-A and III-B effector complexes^13,17,18^. Comparison of the target-bound complex with the binary complex shows only minor conformational changes throughout the Cas7 backbone. However, there are notable rearrangements in the Cas10 subunit. Closer inspection of the two structures reveals an alpha helix (L238-F245) that must be displaced to accommodate the 3’ end of the target RNA strand through the exit channel within Cas10 (Fig 4b). This activation helix appears to communicate long-range allosteric changes, which leads to the opening of the cOA active site cleft. Intriguingly, in the target-bound structure, this cleft can perfectly accommodate a cA_4_ cOA ligand after Csm1 (PDB 6o7B) is aligned to Cas10 in our structure (Fig. 4c). This same analysis of the binary complex shows that this cleft is closed and the cOA has significant steric clashes with the surrounding protein (Fig. 4d). This suggests that the Cas10 subunit of the type III-Dv complex is capable of producing cOA messengers to active Csm6 and other nucleases in a secondary immune response.

**Figure 4:**
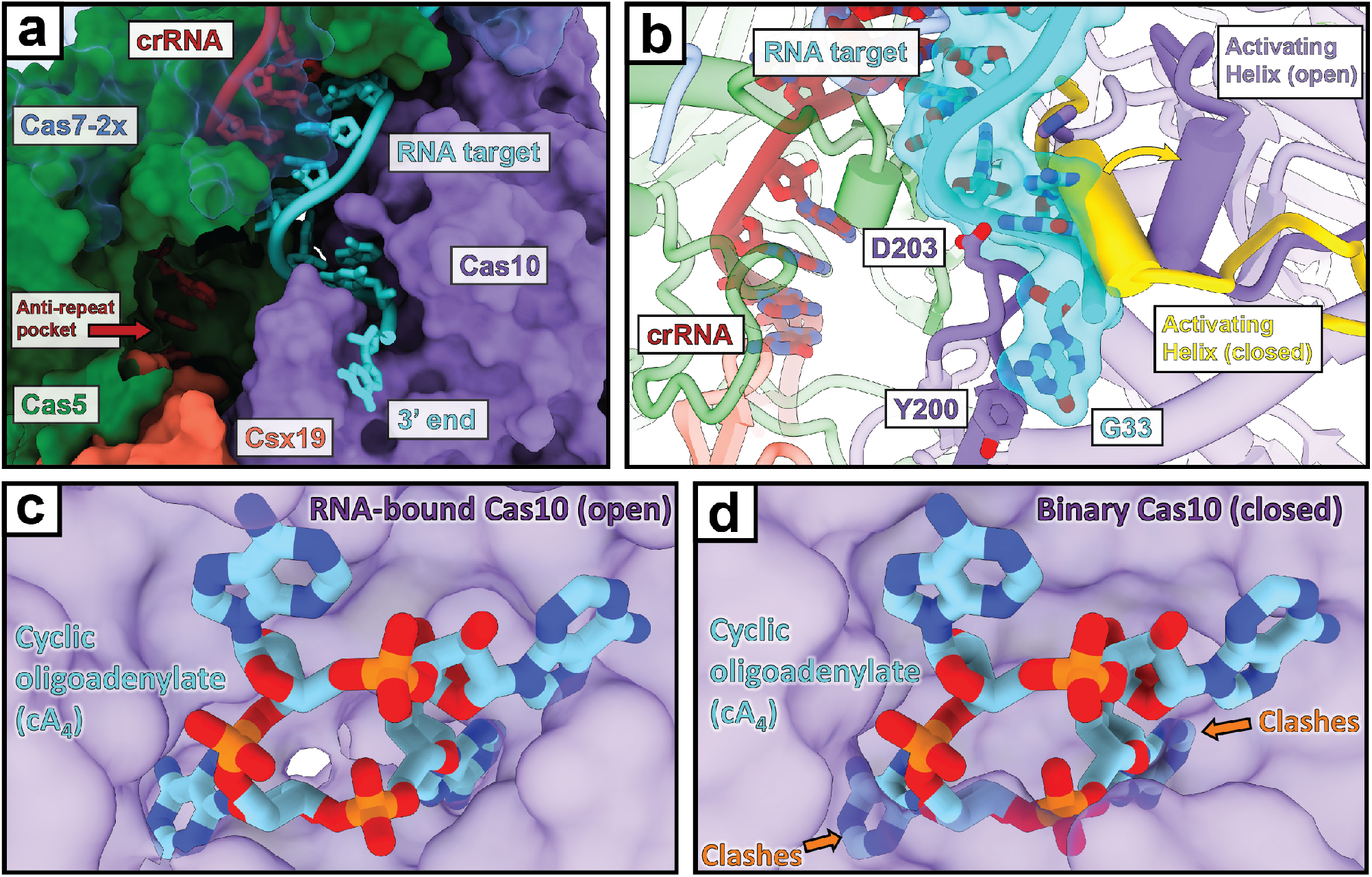
Cas10 activation by non-self RNA binding. **a**, Path of the RNA target strand gets directed through Cas10 rather than into the anti-repeat pocket. **b**, Upon binding a non-self RNA target, an activating helix in Cas10 gets pushed by the target RNA into an active conformation. Yellow arrow indicates directionality of movement of this alpha helix. **c**,**d**, Cyclic oligoadenylate (PDB: 6o7b) bound to the type III-A Cas10 subunit (Csm1) fit into our III-D target-bound and binary models, respectively.

### *In silico* model predictions of the type III-E effector illustrate their evolution from type III-D

To gain a better understanding of the homology between type III-D and III-E systems, we generated an *in silico* atomic model of the *D. ishimotonii* type III-E effector using Alphafold2^26^. In our hypothetical evolutionary progression, III-D1 evolved first and contains single *cas* genes (Supplementary Figure 1). The operon consists of four separate *cas7* genes, *cas10, cas5* (*csx10*), *cas11*, and *csx19*^20,27,28^. Intriguingly, the III-D2 system contains fusions of *cas7* and *cas5* (*cas7-cas5*) and the three following *cas7* genes (*cas7-3x*) and lacks the *csx19* and *cas11* genes. Furthermore, III-D2 has a large domain inserted in the last *cas7* in the operon (*cas7-insertion*). However, type III-E does contain *cas11*, but not *cas5* or *cas10*. These genes are instead fused together into a gene organization of *cas7-cas11-cas7*-*cas7*-(*cas7-insertion*). Strikingly, the III-Dv Cas7-insertion and Cas7.4 of the predicted type III-E model have the protrusion observed from the insertion domain within Cas7, highlighting this as a conserved structural feature between type III-Dv, III-D2, and III-E *cas* operons containing genetic fusions. The type III-E Cas7 backbone follows the same architecture and directionality as our type III-Dv atomic structure, but both differ from the type III-A Csm complex (Fig. 5a,b,c). Özcan and colleagues highlighted two ssRNA cleavage products from the *D. ishimotonii* III-E, which correspond to the active residues of D429 and D654^15^. Interestingly, when aligned to our III-Dv target-bound model, we notice these aspartate residues are positioned right at the scissile phosphodiester bond at positions 31 and 37 of our ssRNA target (Fig. 5d,e). The Cas7 domains that contain these two aspartate residues align with the active site residues D33 and D246 in our Cas7-2x.1 and Cas7-2x.2 domains, respectively. These results paint a clear picture of the conserved structural features between type III-Dv and type III-E. However, despite close alignments of the Cas7 backbone subunits, the type III-E structure is a streamlined version of the type III-Dv complex, with Csx19, Cas5, and Cas10 missing.

**Figure 5:**
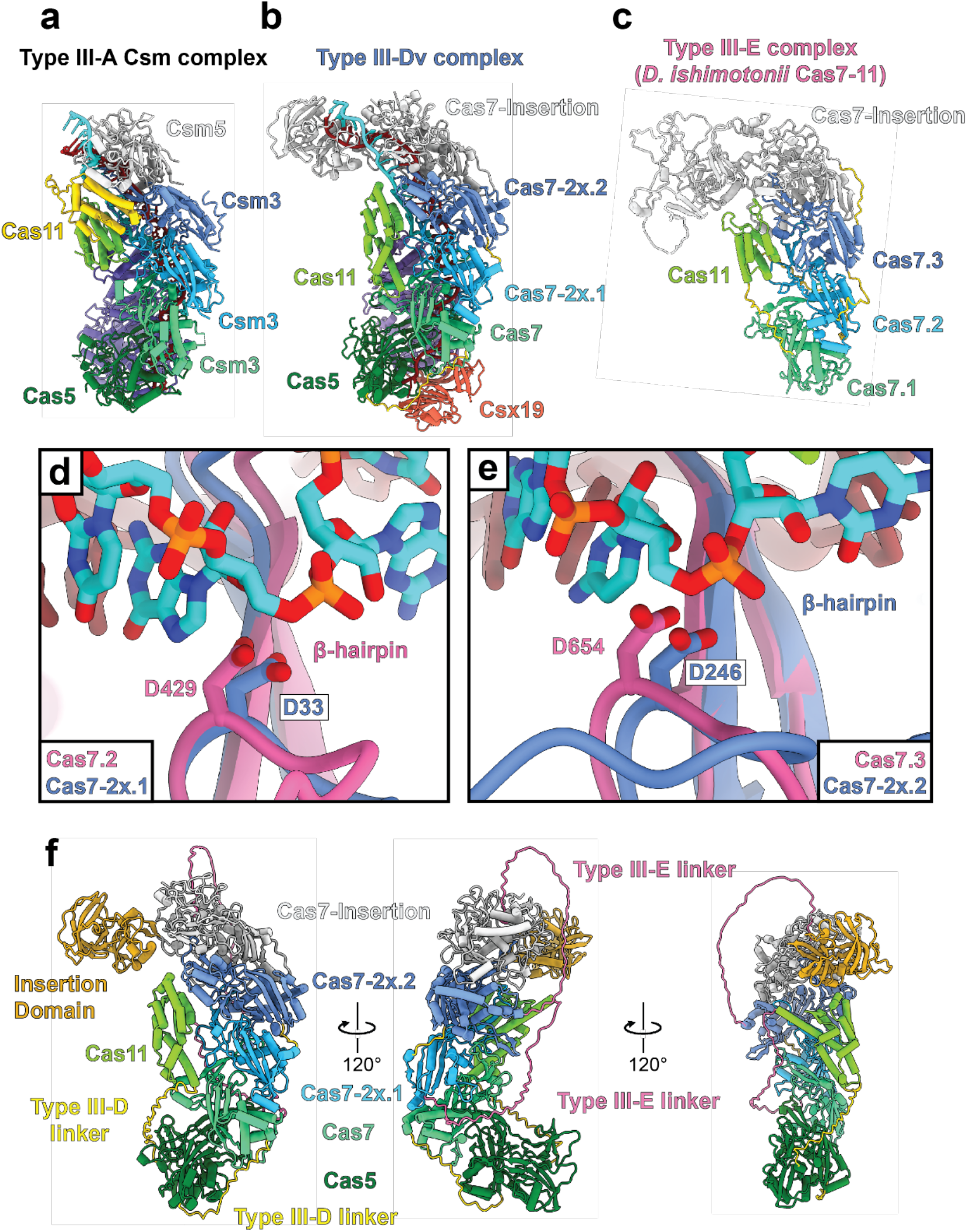
*in silico* model prediction of type III-E and III-D Cas proteins. **a**, Type III-A Csm complex atomic model, colored by subunits. **b**, Type III-Dv atomic model, colored by domains. **c**, Alphafold 2 structure prediction of the *D. ishimotonii* (Cas7-11) type III-E effector protein, colored by domains. Linkers are colored in gold. **d**, Structural alignment of Cas7-2x.1 D33 of type III-Dv with Cas7.2 D429 of *D. ishimotonii* type III-E. **e**, Structural alignment of Cas7-2x.2 D246 of III-Dv with Cas7.3 D654 of *D. ishimotonii* type III-E. **f**, Alphafold 2 predicted models of the a single polypeptide III-Dv containing proteins (Cas7-Cas5-Cas11)-(Cas7-2x)-(Cas7-insertion). Linkers from the III-E model between Cas11 and Cas7-2x, as well as between Cas7-2x and Cas7-insertion are in pink. Linkers in the Cas7-Cas5-Cas11 subunit are in gold.

The fusion of *cas* domains in our III-Dv structure and III-E model involves polypeptide linkers that connect the domains together. We attempted to engineer a single polypeptide type III-Dv effector using subunits from the type III-Dv complex with linkers from type III-E. We linked the C-terminus of the Cas11 domain with the N-terminus of Cas7-2x with the analogous linker in the III-E complex, and the C-terminus of Cas7-2x with the N-terminus of Cas7-insertion with the analogous linker in the III-E complex. We also removed the N-terminal 104 residues – as these residues were not observed in our structures and were necessary for cleavage of an RNA target (Supplementary Fig. 13). The residues in these linkers had no sequence conservation and were only characterized by the presence of flexible amino acids, likely aiding assembly of the domains. We performed Alphafold 2 on the single polypeptide type III-Dv protein lacking Cas10 and Csx19, and remarkably, the structural predictions aligned incredibly well with our III-Dv structure, suggesting that the domains properly fold and assembled together with these long linkers between them (Fig. 5f). This is the first glimpse into an engineered class 1 CRISPR-Cas effector complex into a single polypeptide. This provides an initial blueprint for building user-defined CRISPR-Cas effector complexes for a given activity with the ease of expression and assembly as a single polypeptide.

## Discussion

Recent metagenomic and biochemical analysis of the type III-E single protein effector has led to speculation about how these systems evolved from the multi-subunit type III effectors^15,16,20^. Despite in-depth structural characterization of the type III-B Cmr and type III-A Csm CRISPR-Cas complexes, and a recent structure of the type III-E protein, structures of the type III-D variants are missing. For this reason, our understanding of type III systems is limited to subunit composition and domain organization.

Our structures of the type III-Dv effector complex reveal a number of unique insights. Type III-Dv utilizes fascinating subunit fusions while maintaining the domain organization from type III-D1. Many of the subunits individually align well to the type III-A and type III-B subunits, but the type III-Dv complex does not vary in crRNA length or complex size and is the first in the evolutionary progression of type III systems to utilize subunit fusions. We also discovered a role of the Cas7-insertion domain in RNA target recognition. The insertion domain pulls the crRNA away from the complex at the 3’ end into an alternate geometry, exposing the last six-nts of the crRNA as a seed. Furthermore, we reveal the presence of the unique, uncharacterized subunit Csx19, which nestles between Cas10 and Cas5 at the base of the complex.

Type III CRISPR systems target single-stranded RNA, which makes them powerful post-transcriptional silencers of phage RNA – either mRNA or RNA genomes. Previous studies illustrated a 6-nt ssRNA cleavage periodicity by the III-A, III-B, and III-E effectors^5,6,15^. Here, we show the *in vitro* activity of the III-Dv effector complex on a non-self ssRNA target and reveal three active sites in the Cas7-Cas5-Cas11 and Cas7-2x fusion subunits. Furthermore, we show the absence of cleavage activity by Cas7-insertion. The type III-Dv cleaves on the scale of minutes, emphasizing the potential use of the III-Dv complex as a programmable RNA endonuclease. Rather than recognizing non-self-targets through PAM scanning or PFS requirements, type III complexes rely on recognition of self-transcripts to inhibit activation of Cas10 nuclease activity and production of cOA messengers^10–13,17^. This prevents induction of a secondary immune response where cOAs allosterically activate secondary nucleases for non-specific degradation of nearby nucleic acids. Conformational changes between our structures of the binary and ternary complex of type III-Dv reveal how a non-self ssRNA target activates Cas10 through the displacement of an activating helix at the RNA exit channel of Cas10. These findings elucidate similarities between III-Dv with the previously structurally characterized type III-A Csm and III-B Cmr complexes^12,13,18^.

Overall, our findings highlight the importance of type III-Dv in the evolution of type III CRISPR systems. Previously, we reported on the structure of the *Synechocystis* type I-D complex^29,30^. Our structural work highlighting subunit fusions and insertions in the type III-Dv complex indicate that that I-D and III-Dv likely diverged from a type III-A or III-B ancestor. Type I-D likely served as an intermediate in the rise of type I systems, while III-Dv likely evolved to the single-protein III-E effectors.

The structures of the type III-Dv and III-E effectors provide insights into engineering multi-subunit class 1 complexes into single polypeptide effectors that may be more suitable for biotechnological applications. These systems utilize flexible linkers that hold different domains together, aiding their assembly into a CRISPR-Cas effector. The type III-E operon utilizes these linkers to express a single polypeptide effector, but the type III-Dv system has a mix of subunit fusions and single-gene subunits. We utilized the linkers present in the type III-E complex to design a single polypeptide type III-Dv complex that does not contain Cas10 or Csx19 and found that this design folds nearly identical to our type III-Dv structure, highlighting the future potential of single polypeptide class 1 CRISPR-Cas effector design.

## Methods

Refer to Supplementary Tables 1 and 2 for lists of all plasmids and oligonucleotides, respectively, used in this study.

### Culture conditions

Refer to Supplementary Table 3 for a list of all strains used in this study.

Unless otherwise noted, *Escherichia coli* strains were grown at 37°C in Lysogeny Broth (LB), or on LB-agar (LBA) plates with 1.5% (w/v) agar. Media were supplemented with antibiotics when required as follows: chloramphenicol (Cm; 25 µg/mL), and kanamycin (Km; 50 µg/mL).

### Construction of plasmids

A plasmid (pPF2434) for expression of Cas10, Cas7-5-11, Cas7-2x, Csx19 and Cas7-insert was constructed by PCR-amplifying their genes (primers PF4851+ PF4852) using *Synechocystis* genomic DNA as template and cloning the product into pRSF-1b via KpnI and PstI restriction sites. The *cas10* gene was cloned to incorporate an N-terminal His6 tag followed by TEV protease recognition sequence.

A plasmid (pPF2441) for expression of the first spacer (5’-TGTAGTAGAACCAATCGGGGTCGTCAATAACTCCCG-3’) and flanking repeat sequences (5’-GTTCAACACCCTCTTTTCCCCGTCAGGGGACTGAAAC-3’) from the type III-Dv associated CRISPR array was constructed by PCR-amplifying this region from *Synechocystis* genomic DNA (primers PF4847+ PF4848) and cloning the product into pACYCDuet-1 via NdeI and KpnI restriction sites. A plasmid (pPF2442) was constructed for expression of Cas6-2a with the first spacer and flanking repeat sequences by PCR-amplifying *cas6-2a* (primers PF4849+ PF4850) using *Synechocystis* genomic DNA as template and cloning the product into pPF2441 via NcoI and BamHI restriction sites.

Plasmids pPF3085, pPF3086, pPF3089, pPF3205, and pPF3206 are for expression of mutants Cas7-2x(D29A,D31A,D33A), Cas7-2x(D241A,D246A), ∆Csx19 (nonsense mutation), Cas7-5-11(D26A) and Cas7-insert(∆104 N-terminal residues), respectively. Plasmids pPF3085, pPF3086, pPF3089, pPF3205, and pPF3206 were constructed by site-directed mutagenesis through amplifying plasmid pPF2434 with primers PF5991+PF5992, PF5993+PF5994, PF6281+PF6282, PF6423+PF6424, and PF6425+PF6426, respectively. Each were treated with DpnI to remove PCR template, and Gibson assembly was used to ligate the PCR product into the mutated plasmid.

### Expression and Purification of type III-Dv effector complex

Type III-Dv complex with N-terminal His_6_-TEV-Cas10 was expressed in LOBSTR cells containing plasmids pPF2434 and pPF2442. Five hundred mL cultures were induced with 0.5 mM IPTG at OD_600_ = 0.6 and grown overnight at 18°C. Cells were harvested at 10,000 x g for 10 min. The cell pellet was resuspended in 20 mL of lysis buffer (50 mM HEPES-NaOH, pH 7.5, 300 mM KCl, 5% Glycerol, 1 mM DTT, 10 mM imidazole) supplemented with 0.02 mg/mL DNaseI and cOmplete EDTA free protease inhibitor (Roche). Cells were lysed by a French pressure cell press (American Industry Company) at 10,000 psi, and the lysate was clarified by centrifugation at 15,000 x g for 15 min. The lysate was applied to a HisTrap affinity column (GE Healthcare) equilibrated in lysate buffer and eluted using a gradient against lysate buffer containing 500 mM imidazole. The fractions containing the type III-Dv complex were pooled, treated with TEV protease and incubated at 4°C during overnight dialysis in SEC buffer (10 mM HEPES-NaOH, pH 7.5, 100 mM KCl, 5% Glycerol, 1 mM DTT). The sample was applied to a second HisTrap column; however, due to inefficient TEV cleavage, the complex unexpectedly bound the column and eluted with high imidazole. The complex was further purified by size exclusion chromatography (SEC) on a HiLoad 16/600 Superdex 200 column (GE Healthcare) equilibrated in SEC Buffer. Mutant type III-Dv complexes were similarly expressed and purified, except TEV protease was omitted. Purified complexes were typically concentrated to 1.5 mg/mL using a centrifugal concentrator (Amicon; 100 kDa MWCO), aliquoted, and stored at -80°C.

### Native Mass Spectrometry

5 µL aliquots of the CRISPR complex solution were buffer exchanged into 100 mM ammonium acetate using Biospin P-6 gel columns (Bio-Rad Laboratories Inc., Hercules, CA) prior to native mass spectrometry. Samples were loaded into gold/palladium-coated borosilicate static emitters and subjected to electrospray ionization using a source voltage of 1.0 – 1.3 kV and analyzed in the positive ion mode on a Thermo Scientific Q Exactive Plus UHMR Orbitrap mass spectrometer (Bremen, Germany). Subcomplexes and ejected subunits were produced and measured via quadrupole isolation of the intact complex charge envelope, followed by higher-energy collisional dissociation (HCD) using 290 eV normalized collision energy (NCE). Ion optics and trapping gas pressure were tuned for the transmission and detection of each set of analytes, including the intact complex, subcomplexes, and ejected subunit ions. Native mass spectra were collected by averaging 500 microscans at a resolution of 1,625 at m/z 200. Spectra were deconvoluted using UniDec.

Denaturing liquid chromatography mass spectrometry (LC-MS) was performed on a Dionex UltiMate 3000 nanoLC system coupled to a Thermo Orbitrap Fusion Lumos Tribrid mass spectrometer (San Jose, Ca). The trap column (3 cm) and analytical column (30 cm) were packed in-house with polymer reverse-phase (PLRP) packing material. Approximately 80 ng of the CRISPR complex were injected and subjected to reverse-phase chromatography, utilizing water with 0.1% formic acid as mobile phase A (MPA), and acetonitrile with 0.1% formic acid as mobile phase B (MPB). Forward trapping occurred for 5 minutes at 2% MPB at a flow rate of 5 µL/min at the trap column. Elution onto the analytical column (at 0.3 µL/min) occurred by increasing MPB to 10% over a 3-minute gradient followed by an increase to 35% MPB over 32 minutes. Mass spectra were collected at a resolution of 15,000 at m/z 200, using 5 microscans and an AGC target of 1E6. Spectra were manually averaged over each subunit elution period and deconvoluted with UniDec.

### RNA cleavage by the type III-Dv effector complex

The RNA substrates contained sequence 5’-CAUGACGGAUCGCGGGAGUUAUUGACGACCCCGAUUGGUUCUACUACAAACGUGAUACUA-3’, which included sequence complementary to the type III-Dv crRNA and were fluorescently labelled with either 5’ 6-FAM (PF6575), 5’ IRD800 (PF5856; IRD), or 3’ 6-FAM (PF6576).

RNA cleavage assays were conducted in 5 µL of reaction typically containing 200 nM purified type III-Dv effector complex, 100 nM RNA substrate in final buffer conditions of 6 mM HEPES-NaOH, pH 7.5, 60 mM KCl, 10 mM MgCl_2_ or MnCl_2_, 3% glycerol, 1 mM DTT. Reactions were incubated at 37°C for 30 min or as indicated. Reactions were stopped by adding 1 µL 6 M guanidinium thiocyanate and 12 µL 2x RNA loading dye. Samples were heated for 5 min at 95°C and immediately placed on ice for 3 min. Samples were analyzed by 1x TBE, 15% acrylamide, 8M urea denaturing PAGE (Thermo Fisher). Fluorescent probes were imaged using the Odyssey Fc imaging system (LICOR).

### Cryo-EM grid preparation and data collection

Fully assembled type III-D binary complex was diluted to a concentration of 0.3mg/mL in SEC buffer before 2.5 μL of sample was added to a quantifoil 1.2/1.3 grid that was glow discharged for 1 minute. Sample was applied to the grid in an FEI Vitrobot MarkIV kept at 100% humidity and 4°C before blotting for 5.5 seconds with a force of 0. For the RNA target-bound complex, non-self RNA target (PF5855) was mixed with the binary complex with a 2:1 molar ratio of RNA:Binary complex for 30 minutes at 30° C in SEC buffer to a final protein concentration of 0.3 mg/mL. Grids of the target-bound complex were frozen identical to that of the binary complex. Both grids were loaded to an FEI Titan Krios (Sauer Structural Biology Lab, University of Texas at Austin) operating at 300kV. Images were taken at a pixel size of 0.81 Å/pixel with a dose rate of 10.6 e^-^ /pixel/s for 5 seconds using a Gatan K3 direct electron detector, giving a final dosage of 80.5 e^-^/Å^2^. Data collection was automated using SerialEM using a defocus range of -1.2 to -2.2 µm.

### Cryo-EM data processing

Movies from the Gatan K3 were motion corrected using motioncor2, and corrected micrographs were uploaded to cryoSPARC v2^31^. After CTF correction, initial templates for template-based picking were generated using a blob picker and 2D classification. Template-based particle picking resulted in ∼1.85 million particles (binary complex) and ∼1.92 million particles (target-bound complex) being picked.

Processing the dataset for the binary complex was started with one round of 2D classification, sorting out particles to a new subset of ∼926k particles. Ab initio reconstruction and subsequent heterogeneous refinement with four classes was utilized and ∼649k particles were selected from one of the classes. Particles were then split by exposure groups before performing a final non-uniform (NU) refinement^32^ with per-particle defocus optimization, exposure group CTF parameter optimization, and over per-particle scale minimization. The subunit models were generated using Alphafold 2 and fit into the map using Namdinator and ISOLDE^26,33,34^. The model of the full complex was refined using Phenix real-space-refinement with a non-bonded weight of 250, target bonds rmsd of 0.001, and target angles rmsd of 0.1^35^. The final model yielded from this refinement is composed of ∼649k particles at 2.5 Å resolution. **Data processing workflow can be found in Supplementary Fig. 14**.

For the target-bound complex, ∼1.92 million particles were input into 2D classification and filtering, sorting out particles to a new subset of ∼1.07 million particles. This new subset was then input into ab initio reconstruction and heterogeneous refinement on cryoSPARC v2 with four classes and filtered out ∼453k particles to a new subset of ∼614k particles^31^. These particles were split by exposure groups before performing NU refinement with identical settings to the final NU refinement in the binary complex dataset^32^. The full complex model was refined identical to that of the binary complex. This refinement yielded a 2.8 Å resolution structure from ∼610k particles. **Data processing workflow can be found in Supplementary Fig. 15**.

### *In silico* subunit modeling and refinement

Protein structures of III-D2 Cas7-3x, the Sb-gRAMP III-E effector, the D. ishimotonii III-E effector, and III-D1 Cas7-Cas5 were predicted using Texas Advanced Computing Center Stampede2 computer cluster with AlphaFold2^26^. Structures were predicted using the monomer model preset. The reduced database precision was used for the multiple sequence alignment. The AF2 job run included a relaxation step, resulting in both relaxed and unrelaxed models. Each job was run for a total of 48 hours, yielding 2 to 5 models per protein.

## Acknowledgments

We thank A. Brilot for expert cryo-EM assistance; members of the Taylor, Brodbelt, and Fineran labs for helpful discussions. Data were collected at the Sauer Structural Biology Laboratory at the University of Texas at Austin. This work was supported in part by National Institute of General Medical Sciences (NIGMS) of the National Institutes of Health (NIH) R35GM138348 (to D.W.T.), National Cancer Institute of the NIH F31CA257404 (to L.A.M.) and R35GM139658 (to J.S.B.), Welch Foundation Research Grant F-1938 (to D.W.T.) and F-1155 (to J.S.B.), a Marsden Fund Fast-Start Grant (to R.D.F.) from the Royal Society of New Zealand (RSNZ), the Marsden Fund (to P.C.F.), Bioprotection Aotearoa (Tertiary Education Commission, NZ) (to R.D.F. and P.C.F.). D.W.T is a CPRIT Scholar supported by the Cancer Prevention and Research Institute of Texas (RR160088). The content is solely the responsibility of the authors and does not necessarily represent the official views of the National Institutes of Health.

## Author contributions

R.D.F, P.D.F., and D.W.T. conceived the study. R.D.F., P.D.F., D.W.T., E.A.S., and J.P.K.B. designed the experiments. E.A.S. performed cryo-EM, structure determination, and modeling with assistance from J.P.K.B. L.A.M. performed mass spectrometry. C.L.M. performed in silico structure prediction and comparison. T.L.D. performed initial cleavage experiments. R.D.F. performed cloning, expression, and purification of the complexes and final cleavage experiments. All authors analyzed and interpreted the data. E.A.S., P.C.F., R.D.F. and D.W.T. wrote the manuscript with input from all authors. J.S.B., P.C.F., R.D.F., and D.W.T. supervised and obtained funding for the work.

## References

1. Kato, K. et al. Structure and engineering of the type III-E CRISPR-Cas7-11 effector complex. Cell (2022) doi:10.1016/J.CELL.2022.05.003.

2. Barrangou, R. et al. CRISPR provides acquired resistance against viruses in prokaryotes. Science (1979) 315, 1709–1712 (2007).

3. Hampton, H. G., Watson, B. N. J. & Fineran, P. C. The arms race between bacteria and their phage foes. Nature 577, 327–336 (2020).

4. Brouns, S. J. J. et al. Small CRISPR RNAs guide antiviral defense in prokaryotes. Science (1979) 321, 960–964 (2008).

5. Staals, R. H. J. et al. Structure and activity of the RNA-targeting Type III-B CRISPR-Cas complex of Thermus thermophilus. Mol Cell 52, 135 (2013).

6. Tamulaitis, G. et al. Programmable RNA Shredding by the Type III-A CRISPR-Cas System of Streptococcus thermophilus. Molecular Cell 56, 506–517 (2014).

7. Elmore, J. R. et al. Bipartite recognition of target RNAs activates DNA cleavage by the Type III-B CRISPR–Cas system. Genes & Development 30, 447 (2016).

8. Kazlauskiene, M., Tamulaitis, G., Kostiuk, G., Venclovas, Č. & Siksnys, V. Spatiotemporal Control of Type III-A CRISPR-Cas Immunity: Coupling DNA Degradation with the Target RNA Recognition. Molecular Cell 62, 295–306 (2016).

9. Samai, P. et al. Co-transcriptional DNA and RNA Cleavage during Type III CRISPR-Cas Immunity. Cell 161, 1164–1174 (2015).

10. Niewoehner, O. et al. Type III CRISPR–Cas systems produce cyclic oligoadenylate second messengers. Nature 2017 548:7669 548, 543–548 (2017).

11. Kazlauskiene, M., Kostiuk, G., Venclovas, Č., Tamulaitis, G. & Siksnys, V. A cyclic oligonucleotide signaling pathway in type III CRISPR-Cas systems. Science 357, 605–609 (2017).

12. Jia, N., Jones, R., Sukenick, G. & Patel, D. J. Second Messenger cA4 Formation within the Composite Csm1 Palm Pocket of Type III-A CRISPR-Cas Csm Complex and Its Release Path. Molecular Cell 75, 933-943.e6 (2019).

13. Sofos, N. et al. Structures of the Cmr-β Complex Reveal the Regulation of the Immunity Mechanism of Type III-B CRISPR-Cas. Molecular Cell 79, 741-757.e7 (2020).

14. Makarova, K. S. et al. Evolutionary and functional classification of the CARF domain superfamily, key sensors in prokaryotic antivirus defense. Nucleic Acids Research 48, 8828–8847 (2020).

15. Özcan, A. et al. Programmable RNA targeting with the single-protein CRISPR effector Cas7-11. Nature 2021 597:7878 597, 720–725 (2021).

16. van Beljouw, S. P. B. et al. The gRAMP CRISPR-Cas effector is an RNA endonuclease complexed with a caspase-like peptidase. Science 373, 1349–1353 (2021).

17. You, L. et al. Structure Studies of the CRISPR-Csm Complex Reveal Mechanism of Co-transcriptional Interference. Cell 176, 239–253.e16 (2019).

18. Jia, N. et al. Type III-A CRISPR-Cas Csm Complexes: Assembly, Periodic RNA Cleavage, DNase Activity Regulation, and Autoimmunity. Molecular Cell 73, 264–277.e5 (2019).

19. Osawa, T., Inanaga, H., Sato, C. & Numata, T. Crystal structure of the crispr-cas RNA silencing cmr complex bound to a target analog. Molecular Cell 58, 418–430 (2015).

20. Makarova, K. S. et al. Evolutionary classification of CRISPR–Cas systems: a burst of class 2 and derived variants. Nature Reviews Microbiology 18, 67–83 (2020).

21. Scholz, I., Lange, S. J., Hein, S., Hess, W. R. & Backofen, R. CRISPR-Cas systems in the cyanobacterium Synechocystis sp. PCC6803 exhibit distinct processing pathways involving at least two Cas6 and a Cmr2 protein. PLoS One 8, (2013).

22. Staals, R. H. J. et al. RNA Targeting by the Type III-A CRISPR-Cas Csm Complex of Thermus thermophilus. Molecular Cell 56, 518–530 (2014).

23. Karch, K. R., Snyder, D. T., Harvey, S. R. & Wysocki, V. H. Native Mass Spectrometry: Recent Progress and Remaining Challenges. https://doi.org/10.1146/annurev-biophys-092721-085421 51, 157–179 (2022).

24. Tamara, S., den Boer, M. A. & Heck, A. J. R. High-Resolution Native Mass Spectrometry. Chemical Reviews 122, 7269–7326 (2022).

25. Holm, L. Using Dali for Protein Structure Comparison. Methods Mol Biol 2112, 29–42 (2020).

26. Jumper, J. et al. Highly accurate protein structure prediction with AlphaFold. Nature 2021 596:7873 596, 583–589 (2021).

27. Rouillon, C. et al. Structure of the CRISPR interference complex CSM reveals key similarities with cascade. Mol Cell 52, 124–134 (2013).

28. Rouillon, C., Athukoralage, J. S., Graham, S., Grüschow, S. & White, M. F. Control of cyclic oligoadenylate synthesis in a type III CRISPR system. Elife 7, (2018).

29. McBride, T. M. et al. Diverse CRISPR-Cas Complexes Require Independent Translation of Small and Large Subunits from a Single Gene. Molecular Cell 80, 971--979.e7 (2020).

30. Schwartz, E. A. et al. Structural rearrangements allow nucleic acid discrimination by type I-D Cascade. Nature Communications 2022 13:1 13, 1–11 (2022).

31. Punjani, A., Rubinstein, J. L., Fleet, D. J. & Brubaker, M. A. CryoSPARC: Algorithms for rapid unsupervised cryo-EM structure determination. Nature Methods 14, 290–296 (2017).

32. Punjani, A., Zhang, H. & Fleet, D. J. Non-uniform refinement: adaptive regularization improves single-particle cryo-EM reconstruction. Nature Methods 17, 1214–1221 (2020).

33. Kidmose, R. T. et al. Namdinator - Automatic molecular dynamics flexible fitting of structural models into cryo-EM and crystallography experimental maps. IUCrJ 6, 526–531 (2019).

34. Croll, T. I. ISOLDE: a physically realistic environment for model building into low-resolution electron-density maps. Acta Crystallographica Section D 74, 519–530 (2018).

35. Afonine, P. v et al. Real-space refinement in PHENIX for cryo-EM and crystallography. Acta Crystallographica Section D: Structural Biology 74, 531–544 (2018).

