## Supplementary Information for "Assembly of multi-subunit fusion proteins into the RNA-targeting type III-D CRISPR-Cas effector complex"

**Supplementary Table 1:** Theoretical masses, experimental masses, and % deviation of the type III-Dv complex, subcomplexes, and individual subunits. Experimental masses were recorded via native and LC-mass spectrometry.

| Type III-Dv Complex | Theoretical mass (Da) | Native MS Experimental mass (Da) | Denaturing LC-MS Experimental mass (Da) | Deviation from expected mass (%)* |
|---|---|---|---|---|
| Cas7_insert•Cas7-Cas5-Cas11•Cas10-His6•Cas7-2x•Csx19•mature crRNA | 331,405 | 332,300 | - | 0.27 |
| Cas7_insert•Cas7-Cas5-Cas11•Cas10-His6•Cas7-2x•mature crRNA | 310,213 | 310,980 | - | 0.25 |
| Cas7_insert•Cas7-Cas5-Cas11•Cas10-His6•Csx19•mature crRNA | 274,778 | 274,840 | - | 0.02 |
| Cas7_insert•Cas7-Cas5-Cas11•Cas7-2x•Csx19•mature crRNA | 267,286 | 267,770 | - | 0.18 |
| Cas10-His6•mature crRNA | 76,055 | - | 76,110 | 0.07 |
| Cas7_insert | 90,080 | - | 90,080 | 0.00 |
| Cas7-Cas5-Cas11 | 87,451 | - | 87,450 | 0.00 |
| Cas10-His6 | 64,119 | 64,170 | 64,120 | 0.00 |
| Cas7-2x | 56,627 | - | 56,630 | 0.01 |
| Csx19 | 21,192 | 21,190 | 21,190 | 0.01 |
| Mature crRNA | 11,936 | - | - | - |

**Supplementary Table 2:** Plasmids used in this study.

| Plasmid | Description | Reference |
|---|---|---|
| pACYCDuet-1 | Two T7/LacO promoters with P15A replicon, $Cm^R$ | Novagen |
| pPF2434 | N-$His_6$-tagged Cas10, Cas7-5-11, Cas7_2x, Csx19 and Cas7-insert, pRSF-1b | This study |
| pPF2441 | Spacer1 of Synechocystis type III-Dv CRISPR array, pACYCDuet-1 | This study |
| pPF2442 | Plasmid pPF2434 with Cas6-2a | This study |
| pPF3085 | Modified pPF2434 with Cas7-2x(D29A,D31A,D33A) | This study |
| pPF3086 | Modified pPF2434 with Cas7-2x(D241A,D246A) | This study |
| pPF3089 | Modified pPF2434 without Csx19 | This study |
| pPF3205 | Modified pPF2434 with Cas7-5-11(D26A) | This study |
| pPF3206 | Modified pPF2434 with Cas7-insert(Δ104 N-terminus) | This study |
| pRSF-1b | T7/LacO promoter with RSF1030-derived replicon, $Km^R$ | Novagen |

**Supplementary Table 3:** Oligonucleotides used in this study.

| Name | Sequence (5'-3') | Notes | Restriction site |
|---|---|---|---|
| PF4847 | TATACATATGGCATACAGACTGTTTTTCAGTGTGATAG | F repeats and spacer1 | NdeI |
| PF4848 | CGAGGGTACCGGGACTCCAACCCCCCAAG | R repeats and spacer1 | KpnI |
| PF4849 | TATACCATGGTGGATCTAAAATCCTTAGCTG | F *cas6-2a* | NcoI |
| PF4850 | ATTCGGATCCTTATTGAACATTGGCTAAGGC | R *cas6-2a* | BamHI |
| PF4851 | TGGGTACCGAAAACCTGTATTTTCAGGGCTTTCTAGTTCTAATTGAGA CTTCCGGTAATC | F *cas10* | KpnI |
| PF4852 | CGGCCGCAAGCTTGTCGACCTGCAGTTAACTAGGTTTGATTGGAAAA CTCTGG | R *cas7-insert* | PstI |
| PF5855 | rCrArUrGrArCrGrGrArUrCrGrCrGrGrArGrUrUrArUrUrGrArCrGrArCrCrC rCrGrArUrUrGrGrUrUrCrUrArCrUrArCrArArArCrGrUrGrArUrArCrUrA | 60nt RNA target | - |
| PF5991 | GGCGCCGCTGCGACGGCTTTAGCCCTGGCGGTTAATGGTG | F *cas7-2x* D33A mutant | - |
| PF5992 | AAAGCCGTCGCAGCGGCGCCACCCACACCACCAATG | R *cas7-2x* D33A mutant | - |
| PF5993 | GGCTGGACTGGCGATCGCTATTTTGCCCCTCGTTAGTCAAGTG | F *cas7-2x* D246A mutant | - |
| PF5994 | TAGCGATCGCCAGTCCAGCCCCTTCAGCTTTCACCATGAC | R *cas7-2x* D246A mutant | - |
| PF6281 | GCCTAAGTTAGTAACTTTACCACTACCACCAATATGAAATTACCCTC | F Δ*csx19* mutant | - |
| PF6282 | GTAAAGTTACTAACTTAGGCGGCCTCCTGCTG | R Δ *csx19* mutant | - |
| PF6423 | GAACTAGCCAGTGTTGTACAACGGGATGGAG | F *cas7-5-11* D26A mutant | - |
| PF6424 | TGTACAACACTGGCTAGTTCCCCCCGACCCATG | R *cas7-5-11* D26A mutant | - |
| PF6425 | AAGTAAAATGACACCCAGAAATGTTAACGCTAGCAAC | F *cas7-insert* Δ104 N-term mutant | - |
| PF6426 | TTCTGGGTGTCATTTTACTTAACCTCCAATTTAATTAAACGTTC | R *cas7-insert* Δ104 N-term mutant | - |

| Name | Sequence (5'-3') | Notes | Restriction site |
|---|---|---|---|
| PF5856 | /5IRD800CWN/rCrArUrGrArCrGrGrArUrCrGrCrGrGrGrArGrUrUrArUrUrGrArCrGrArCrCrCrCrGrArUrUrGrGrUrUrCrUrArCrUrArCrArArArCrGrUrGrArUrArCrUrA | 5'-IRD800 60nt RNA target | - |
| PF6575 | /56-FAM/rCrArUrGrArCrGrGrArUrCrGrCrGrGrGrArGrUrUrArUrUrGrArCrGrArCrCrCrCrGrArUrUrGrGrUrUrCrUrArCrUrArCrArArArCrGrUrGrArUrArCrUrA | 5'-FAM 60nt RNA target | - |
| PF6576 | rCrArUrGrArCrGrGrArUrCrGrCrGrGrGrArGrUrUrArUrUrGrArCrGrArCrCrCrCrGrArUrUrGrGrUrUrCrUrArCrUrArCrArArArCrGrUrGrArUrArCrUrA/36-FAM/ | 3'-FAM 60nt RNA target | - |
| PF6577 | /56-FAM/rCrArUrGrArCrGrGrArUrCrGrCrGrGrGrArGrUrUrArUrUrGrArCrGrArCrCrCrCrGrArUrUrGrGrUrUrCrUrA | 5'-FAM 43nt RNA target | - |
| PF6578 | /56-FAM/rCrArUrGrArCrGrGrArUrCrGrCrGrGrGrArGrUrUrArUrUrGrArCrGrArCrCrCrCrGrArUrUrG | 5'-FAM 37nt RNA target | - |
| PF6579 | /56-FAM/rCrArUrGrArCrGrGrArUrCrGrCrGrGrGrArGrUrUrArUrUrGrArCrGrArCrCrC | 5'-FAM 31nt RNA target | - |
| PF6580 | /56-FAM/rCrArUrGrArCrGrGrArUrCrGrCrGrGrGrArGrUrUrArUrUrGrArCrG | 5'-FAM 27nt RNA target | - |
| PF6582 | /56-FAM/rCrArUrGrArCrGrGrArUrCrGrCrGrGrGrArGrUrUrArUrUrGrArCrGrArCrCrCrCrGrArUrUrGrGrUrUrCrUrArCrUrArCrArGrUrUrUrCrArGrUrCrCrCrC | 5'-FAM 60nt RNA anti-repeat | - |
| PF6583 | /5IRD800CWN/rCrArUrGrArCrGrGrArUrCrGrCrGrGrGrArGrUrUrArUrUrGrArCrGrArCrCrCrCrGrArUrUrG | 5'-IRD800 37nt RNA target | - |

**Supplementary Table 4:** Bacterial strains used in this study.

| **Strain** | **Genotype/Phenotype/Description** | **Reference** |
|---|---|---|
| DH5α | *E. coli* $F^-$, ϕ80d*lacZ*ΔM15, Δ(*lacZYA–argF*)U169, *endA1*, *recA1*, *hsd*R17 ($r_K^- m_K^+$), *deoR*, *thi-1*, *sup*E44, $\lambda^-$, *gyr*A96, *rel*A1 | Gibco/BRL |
| LOBSTR | *E. coli* B $F^-$ *ompT, gal, dcm, lon, hsdS*$_B$($r_B^- m_B^-$), λ(DE3 [*lacI lacUV5-T7p07 ind1 sam7 nin5*]) [*malB*$^+$]$_{K-12}$($\lambda^S$) *arnA slyD* | Kerafast |
| *Synechocystis* sp. PCC 6803 | Glucose tolerant laboratory wild-type strain GT-01 | (Morris et al., 2014), (Williams, 1988) |

**Supplementary Table 5:** Model Statistics for the type III-Dv binary and ternary complex.

| | III-Dv Binary Complex | III-Dv Ternary Complex |
|---|---|---|
| **Data collection and processing** | | |
| Magnification | 29000 | 29000 |
| Voltage (kV) | 300 | 300 |
| Electron exposure (e–/Å$^2$) | 80.5 | 80.5 |
| Defocus range (μm) | 1.2 to 2.2 | 1.2 to 2.2 |
| Pixel size (Å) | 0.81 | 0.81 |
| Symmetry imposed | C1 | C1 |
| Initial particle images (no.) | 1,890,840 | 1,919,796 |
| Final particle images (no.) | 648,782 | 609,722 |
| Map resolution (Å)<br>FSC threshold | 2.5<br>0.143 | 2.8<br>0.143 |
| Map resolution range (Å) | N/A | N/A |
| **Refinement** | | |
| Model resolution (Å)<br>FSC threshold | 2.7<br>0.5 | 2.9<br>0.5 |
| Model resolution range (Å) | N/A | N/A |
| Map sharpening *B* factor (Å$^2$) | 99.3 | 124.7 |
| Model composition<br>Non-hydrogen atoms<br>Protein residues<br>Ligands | <br>22168<br>2718<br>0 | <br>22688<br>2697<br>0 |
| *B* factors (Å$^2$)<br>Protein<br>Nucleotide | <br>20.27/98.25/49.61<br>28.70/93.35/46.60 | <br>9.98/84.88/40.98<br>23.07/94.13/37.97 |
| R.m.s. deviations<br>Bond lengths (Å)<br>Bond angles (°) | <br>0.012<br>1.888 | <br>0.013<br>1.947 |
| **Validation**<br>MolProbity score<br>Clashscore<br>Poor rotamers (%) | <br>1.00<br>0.23<br>1.70 | <br>1.24<br>0.34<br>2.69 |
| Ramachandran plot<br>Favored (%)<br>Allowed (%)<br>Disallowed (%) | <br>96.41<br>3.48<br>0.11 | <br>95.82<br>3.85<br>0.34 |

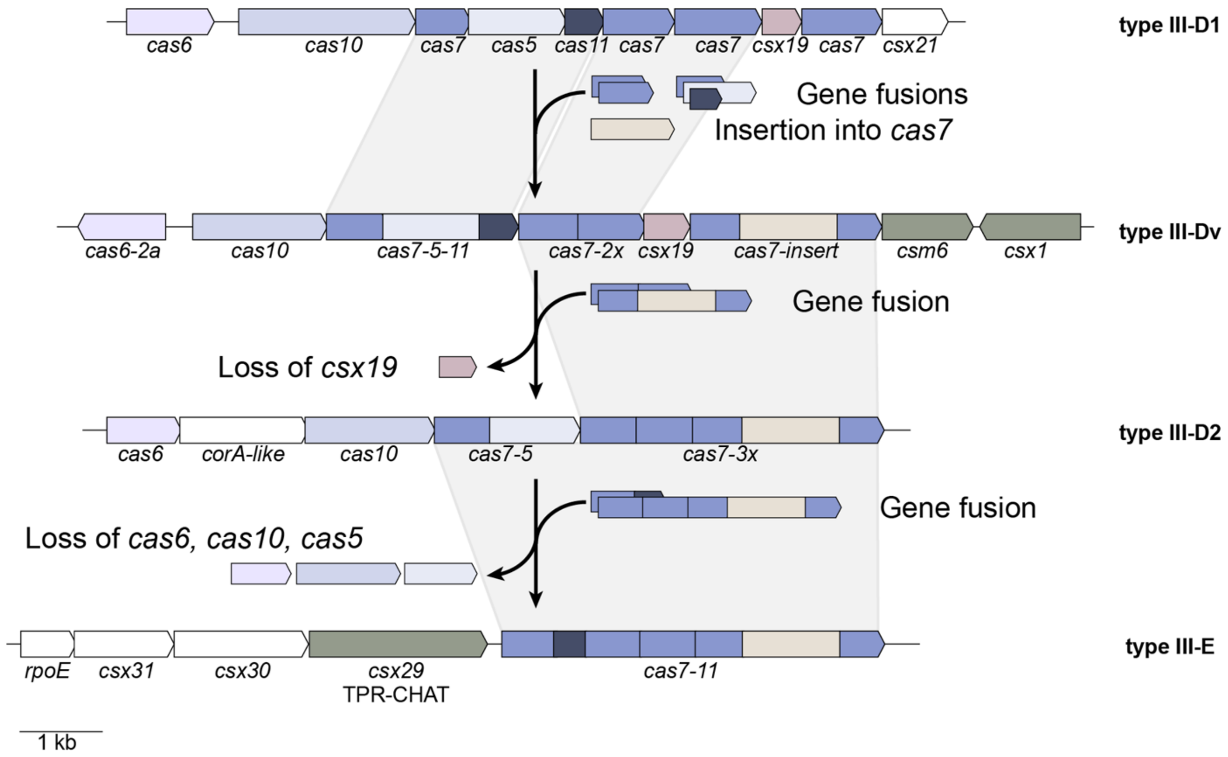


**Supplementary Figure 1: Proposed evolution between type III-D variants to type III-E.**

**a**

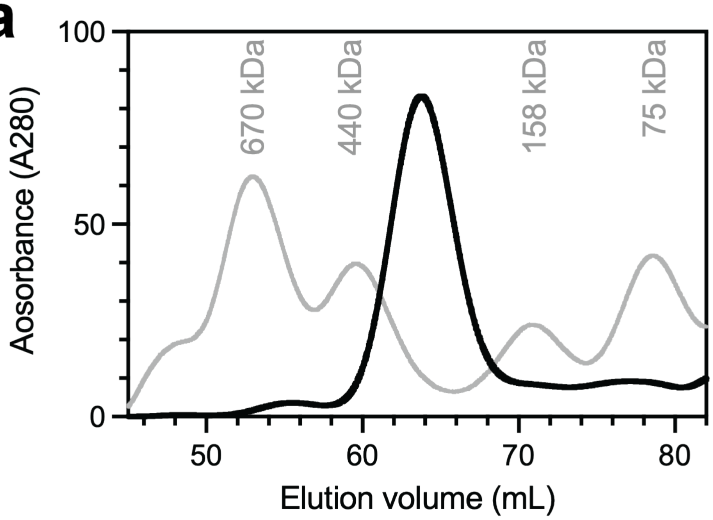


**Supplementary Figure 2: Purification of the type III-Dv effector complex. a,** Size-exclusion chromatograph of the WT type III-Dv complex bound to crRNA. Black peak corresponds to ~330 kDa. Grey peaks represent standardized molecular weights.

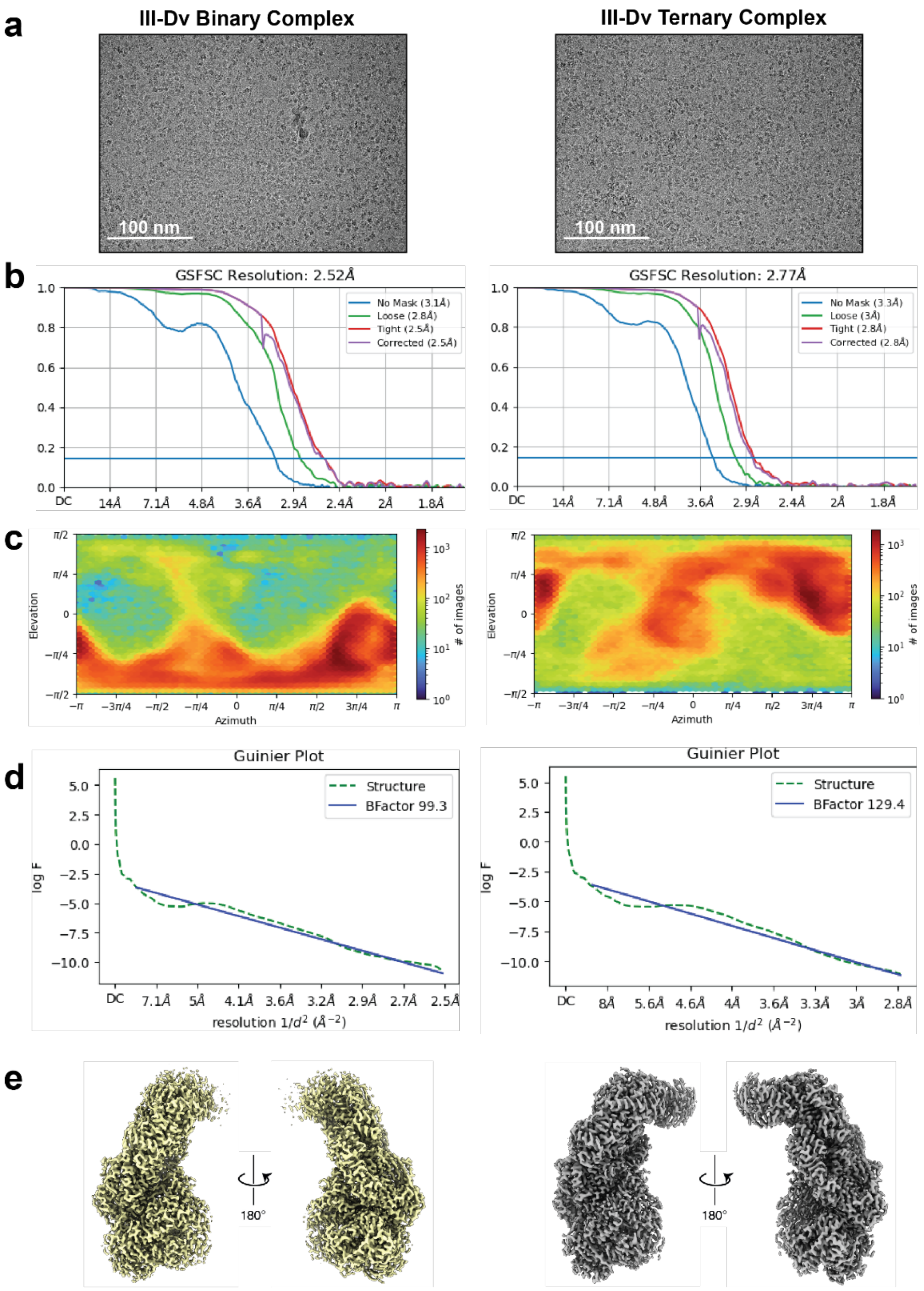


**Supplementary Figure 3: EM validation of type III-Dv binary (left) and ternary (right) maps.** **a,** Representative micrographs. Scale bar is shown at 100 nm. **b,** FSC plot of the two maps based on the 0.143 gold standard of two half maps. **c,** Euler angular distribution showing distribution of orientations that contribute to the final map. **d,** Guinier plots to show BFactor sharpening calculation for final EM maps. **e,** Final sharpened EM maps from cryoSPARC v2 at 2.52 Å and 2.77 Å, respectively.

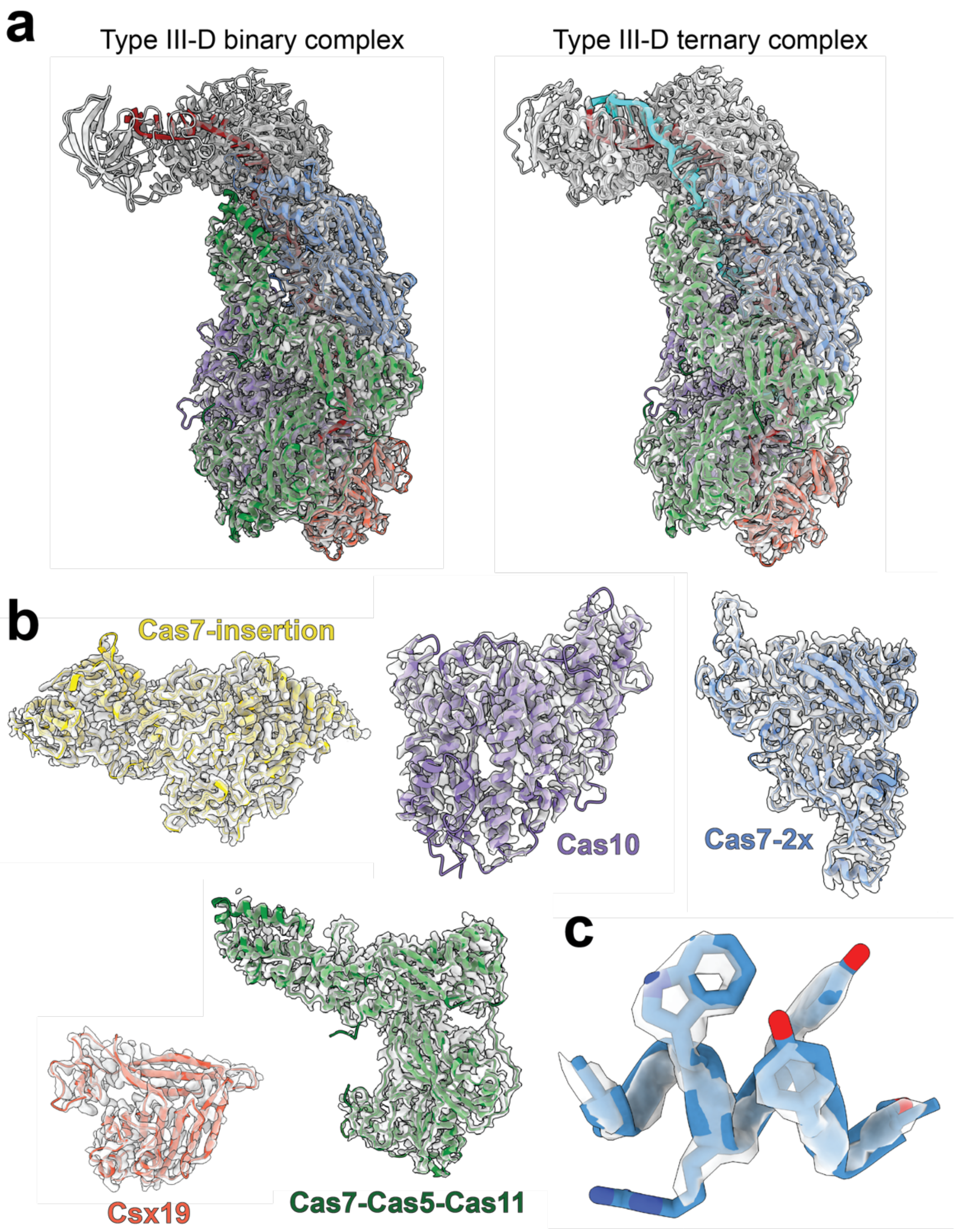


**Supplementary Figure 4: Representative cryo-EM density of the type III-Dv complex. a,** Overall complexes fit into the binary and ternary cryo-EM maps prior to local refinement. **b,** Individual subunits fit into their respective cryo-EM density. **c,** Density of an alpha-helix in the Cas7-insertion subunit.

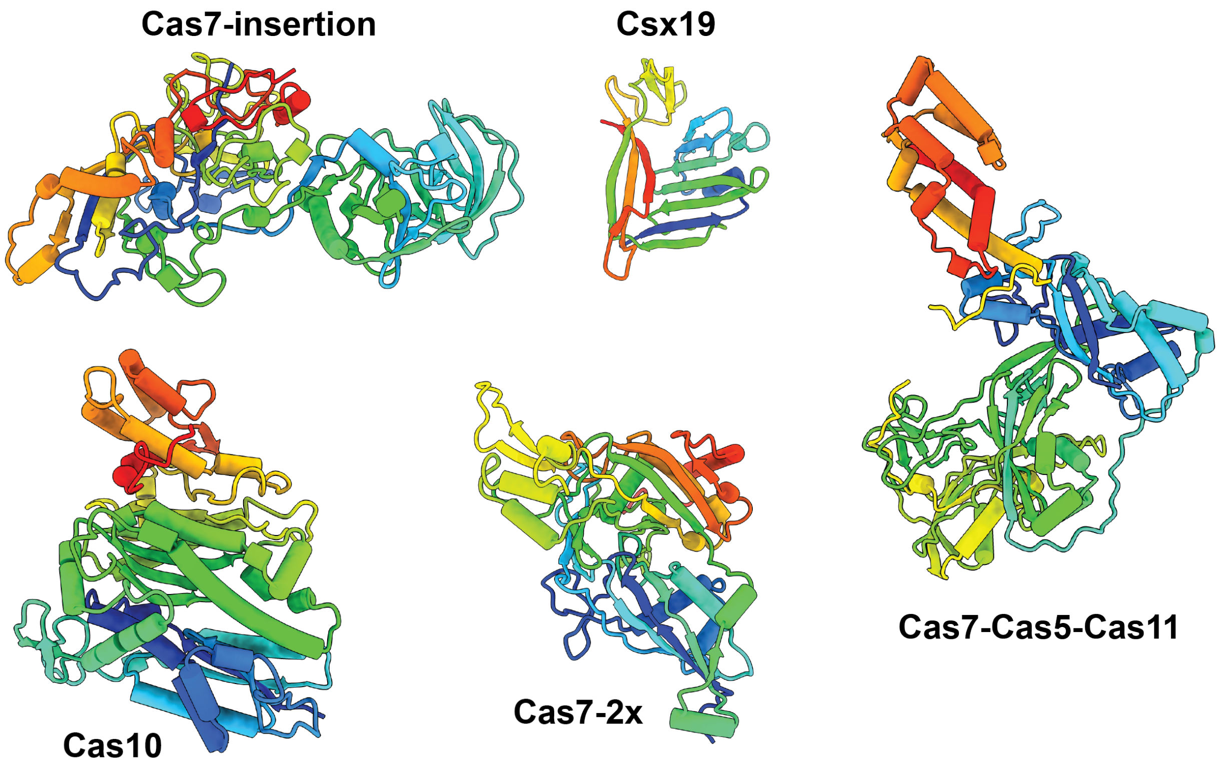


**Supplementary Figure 5: Connectivity of the type III-Dv subunit structures.** The subunits are colored N-terminus (blue) to C-terminus (red).

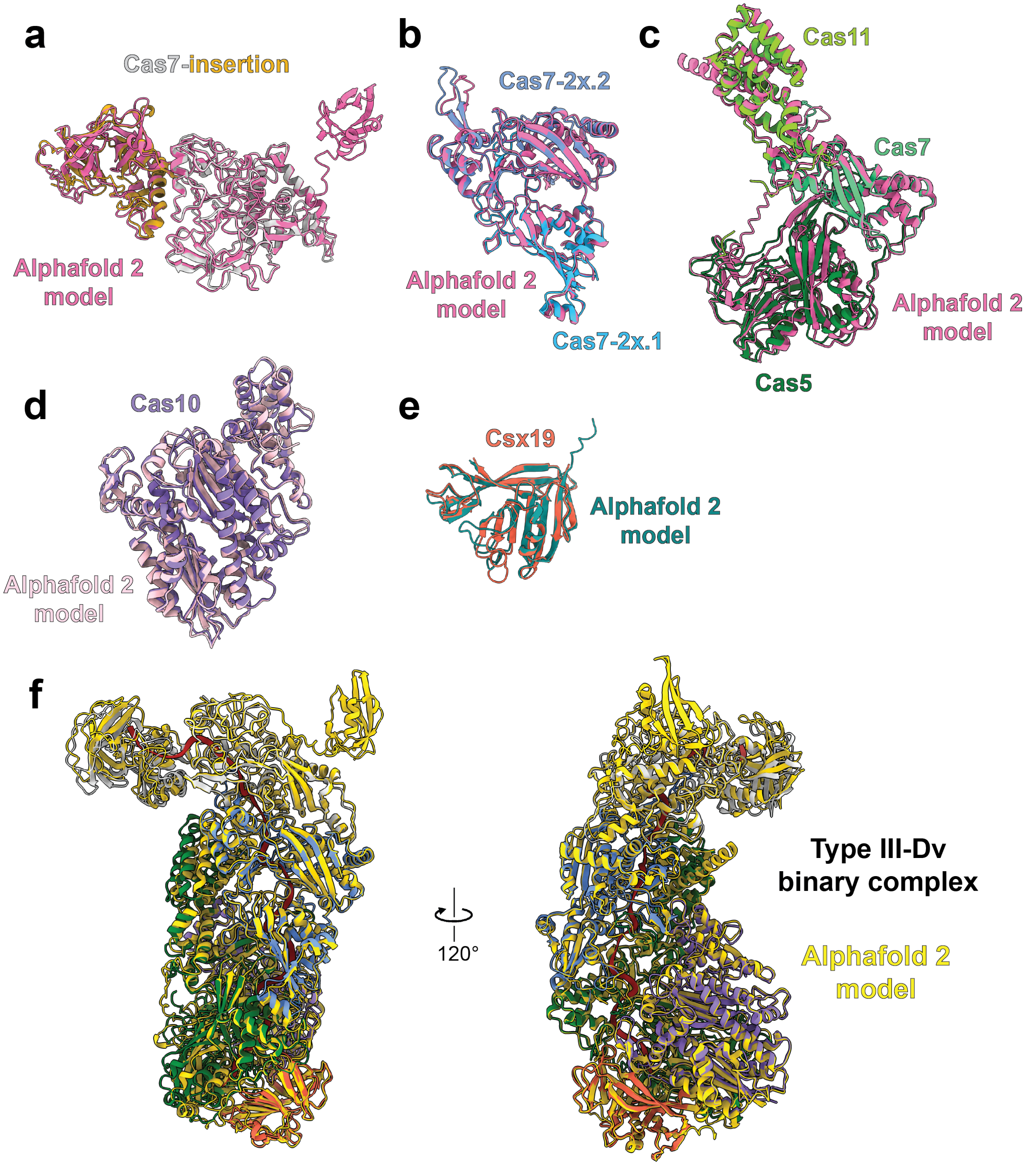


**Supplementary Figure 6: Structural comparisons of Alphafold 2 predictions the final models of a,** Cas7-insertion (RMSD 0.667 Å) ; **b,** Cas7-2x (RMSD 0.915 Å); **c,** Cas7-Cas5-Cas11 (RMSD 1.014 Å); **d,** Cas10 (RMSD 0.971 Å); **e,** Csx19 (RMSD 0.811 Å). All RMSDs were calculated for pruned atom pairs. **f,** Overlay of the Alphafold 2 models onto the final models of their respective subunits assembled into the full binary complex.

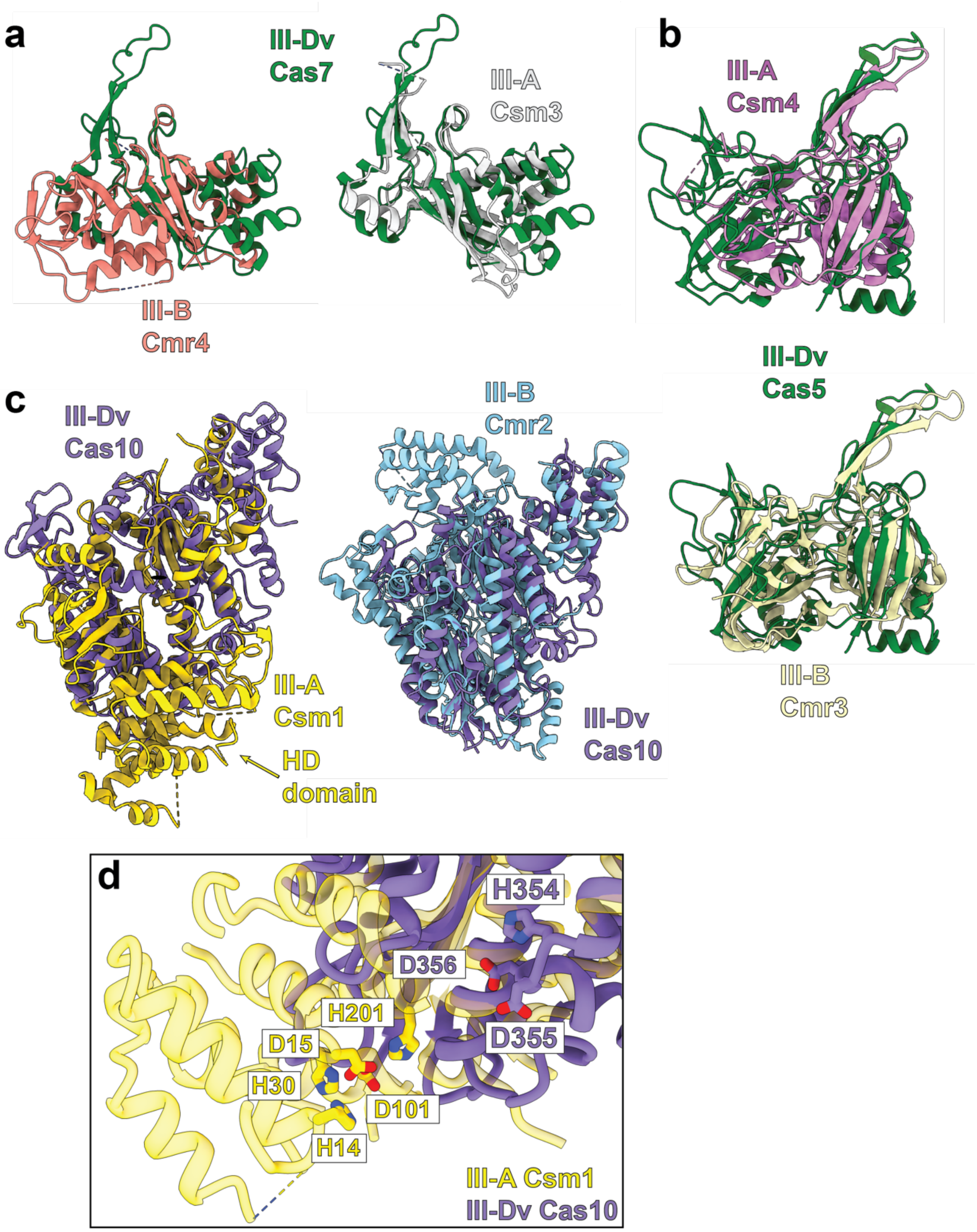


**Supplementary Figure 7: Subunit comparisons between type III-Dv and type III-A/B homologues. a,** crRNA-target duplex of the III-Dv ternary complex is stabilized by a positive patch on the Cas11 domain of Cas7-Cas5-Cas11. **a,** Structural alignments between the III-Dv Cas7 domain of Cas7-Cas5-Cas11 with Cmr4 (Cas7) of the type III-B (peach) and Csm3 of the type III-A complex (white); **b,** III-Dv Cas5 domain of Cas7-Cas5-Cas11 with Cas5 (Csm4) of the type III-A complex (magenta) and Cas5 (Cmr3) of the type III-B complex (beige); **c,** III-Dv Cas10 subunit with Cas10 (Csm1) of the type III-A complex (yellow) and Cas10 (Cmr2) of the type III-B complex (cyan). **d,** HD domain comparison between Csm1 (Cas10) of type III-A with the putative III-Dv HD site.

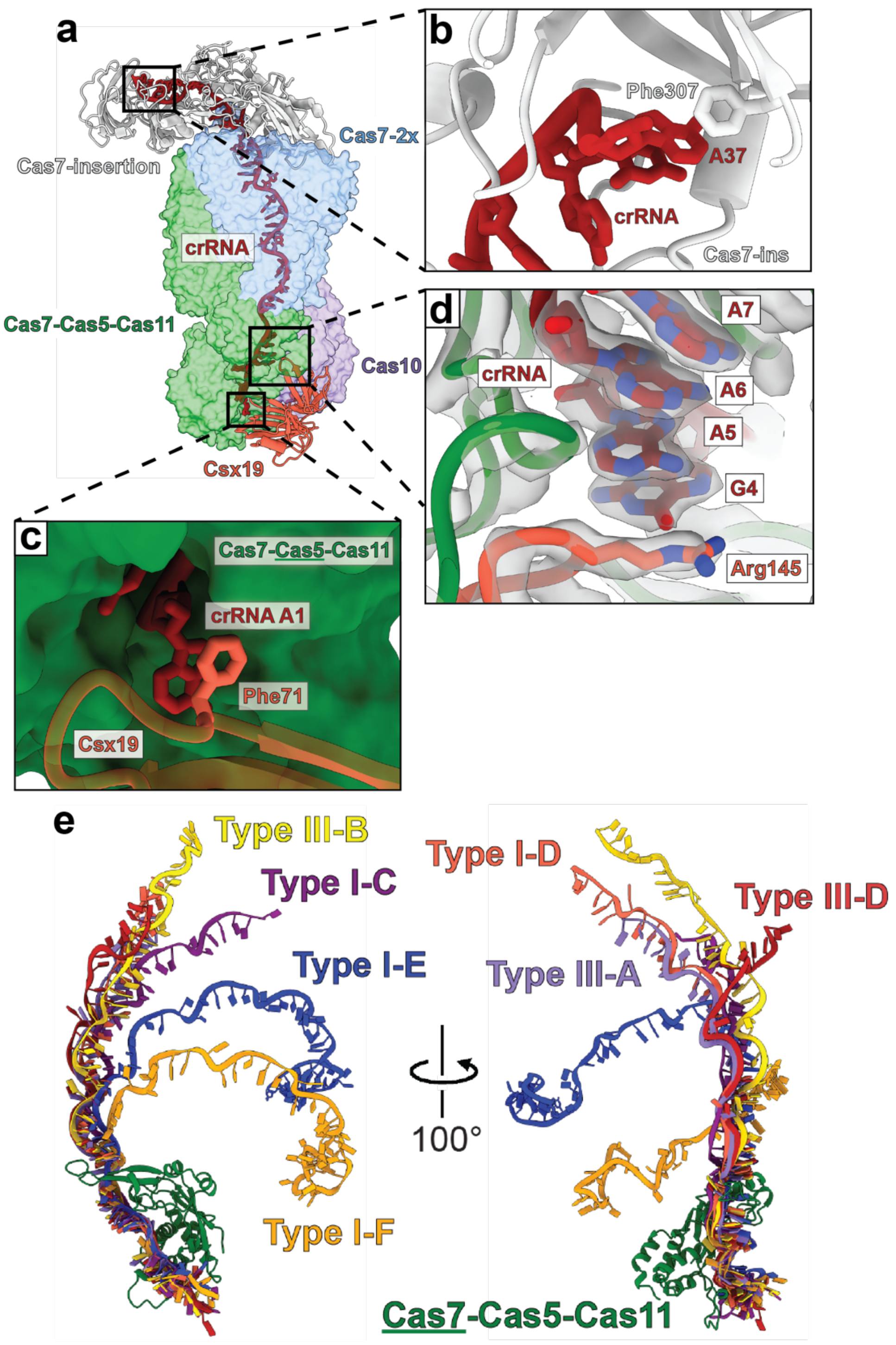


**Supplementary Figure 8: type III-Dv interactions with the crRNA. a,** crRNA trajectory through the type III-Dv effector complex. **b,** 3' crRNA capping by Phe307 of Cas7-insertion. **c,** 5' crRNA capping by Phe71 of Csx19. **d,** Arg145 of Csx19 interacting with G4 of the crRNA, upstream of the 5' crRNA handle. **e,** crRNA geometry comparison between type III-Dv and other type I and type III systems.

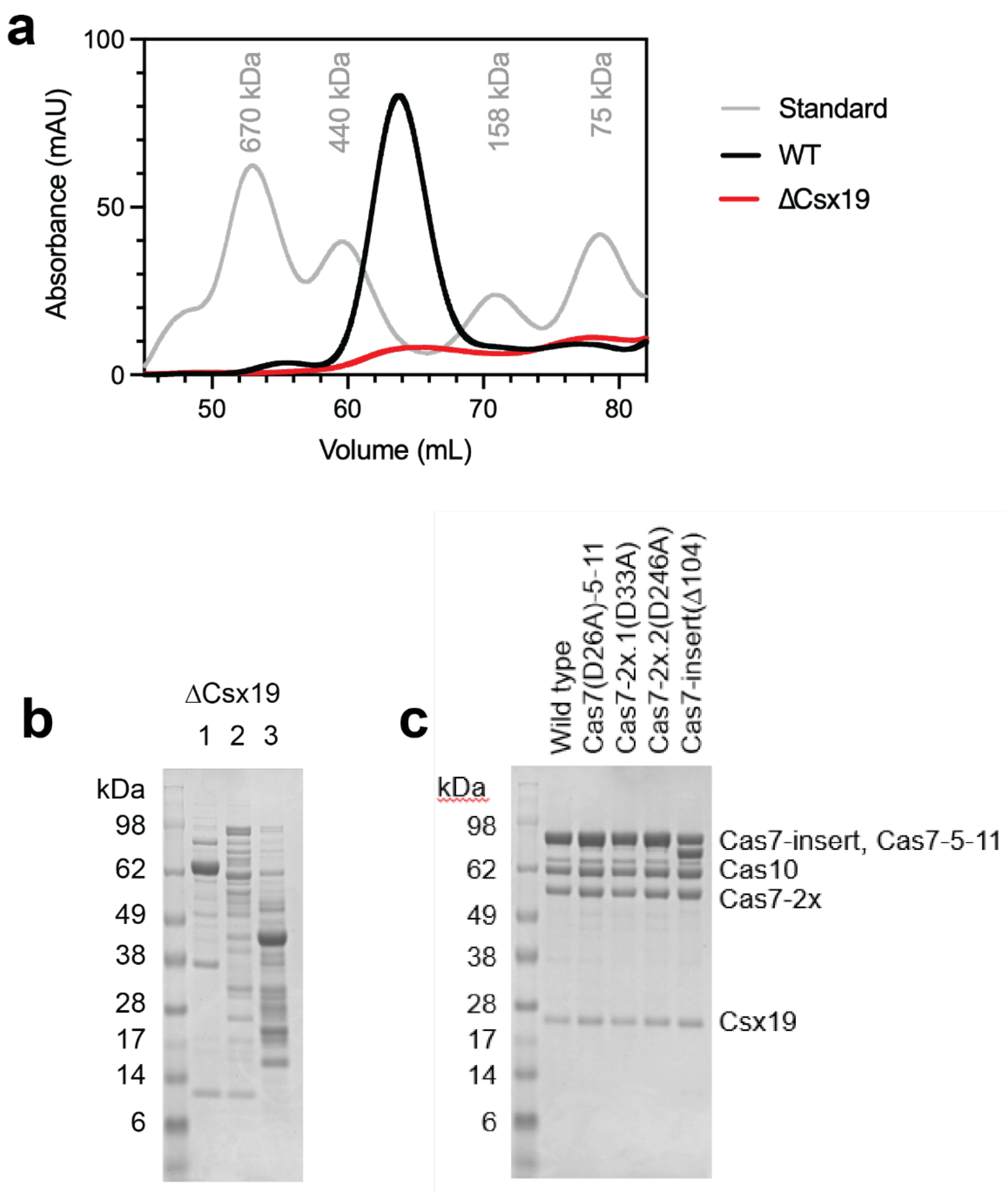


**Supplementary Figure 9: Purification of mutant III-Dv effector complexes. a,** Size-exclusion chromatograph of the ΔCsx19 type III-Dv complex bound to crRNA (red trace). Black peak corresponds to wild-type III-Dv complex. Grey peaks represent standardized molecular weights. **b,** SDS-PAGE of the ΔCsx19 from the two broad peaks seen in **a**. No complex appears to form. **c,** Purification of Cas7 active site mutants for RNA cleavage analysis.

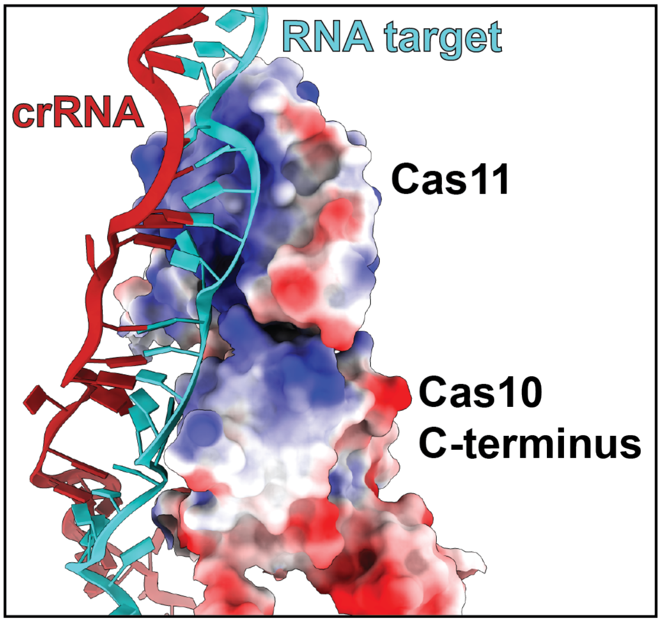


**Supplementary Figure 10: Cas11 interactions with the crRNA.** crRNA-target duplex of the III-Dv ternary complex is stabilized by a positive patch on the Cas11 domain of Cas7-Cas5-Cas11 and the c-terminal domain of Cas10.

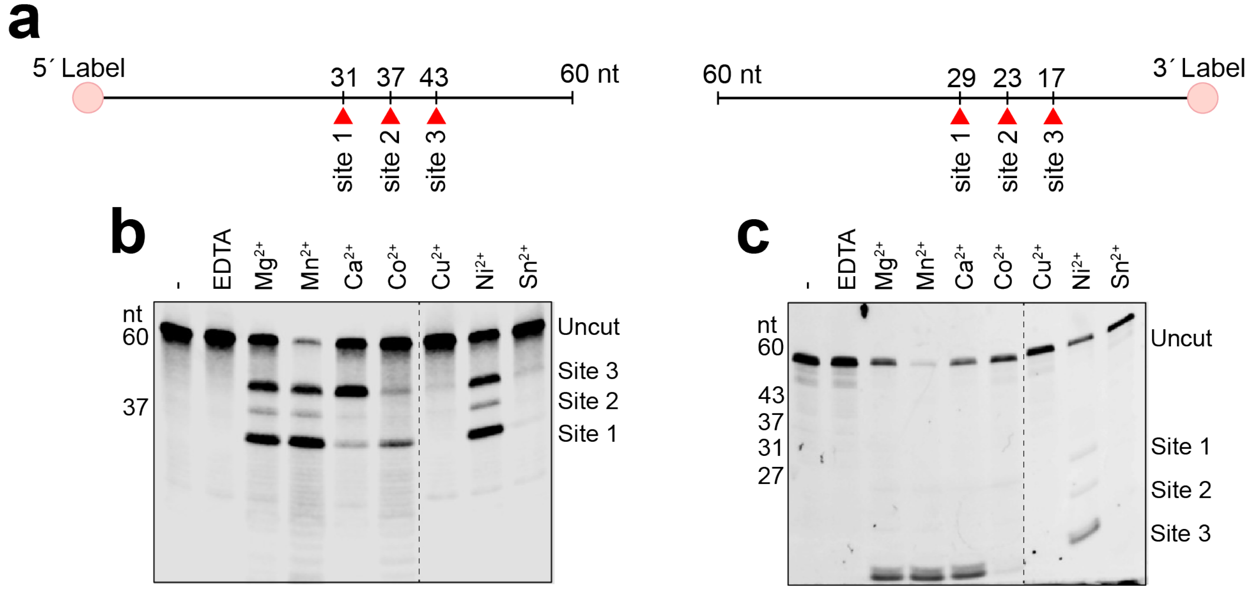


**Supplementary Figure 11: Metal-dependent cleavage of an RNA target by type III-Dv. a,** Fluorescent probes used in tests of cleavage conditions using different divalent cations. **b,** Metal dependent cleavage of the 5'-labelled RNA target. Cleavage site 1, 2, and 3 correspond to 31, 37, and 43 nucleotide products, respectively. **c,** Metal dependent cleavage of the 3'-labelled RNA target. Cleavage site 1, 2, and 3 correspond to 29, 23, and 17 nucleotide products, respectively.

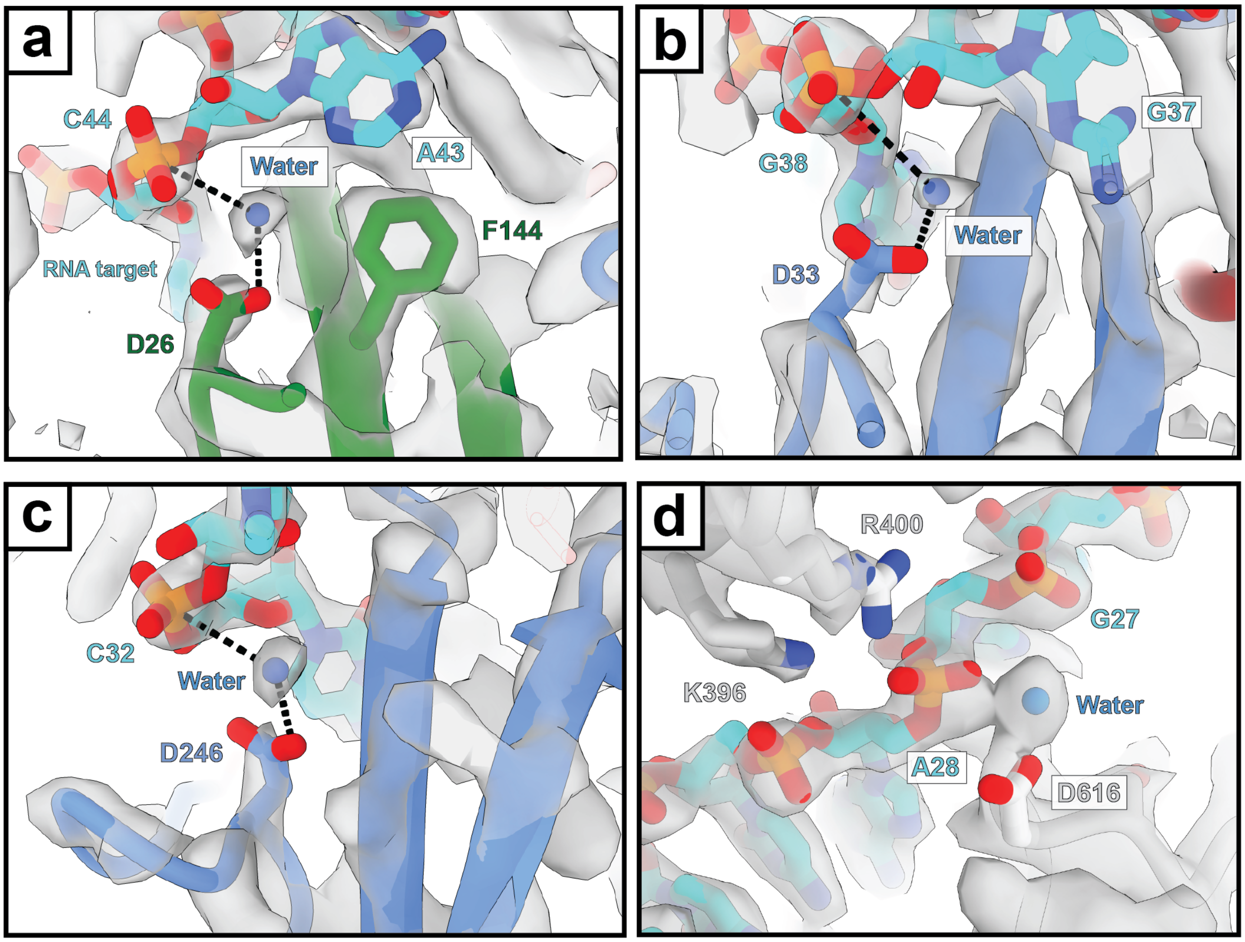


**Supplementary Figure 12: Cas7 domain active sites in the type III-Dv complex. a,** Cas7-Cas5-Cas11 (D26), **b,** Cas7-2x.1 (D33), and **c,** Cas7-2x.2 (D246) active sites with EM density. **d,** Cas7-insertion also coordinates a water molecule, but is not active in cleaving the RNA.

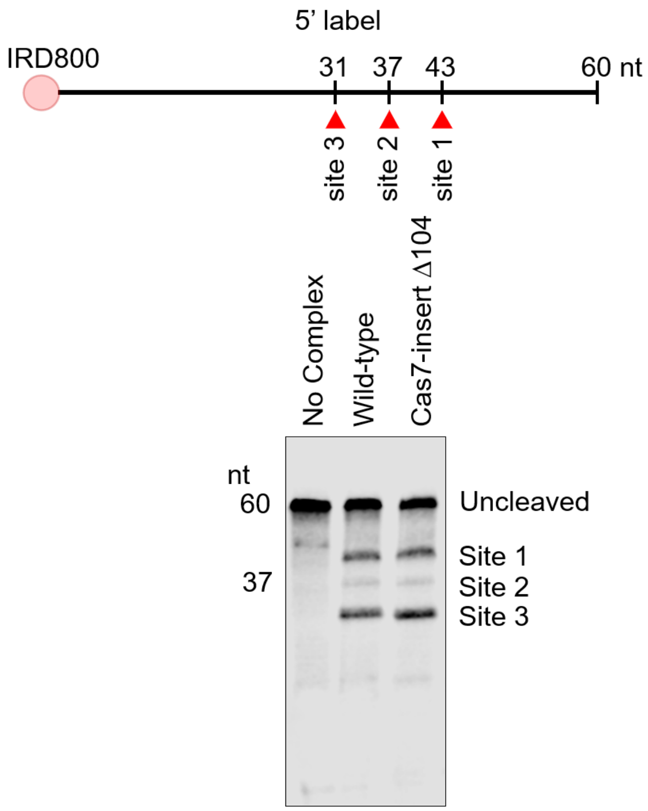


**Supplementary Figure 13: RNA targeting by the Δ104 Cas7-insertion type III-Dv complex.** Site 1, 2, and 3 correspond to cleavage by Cas7-2x.2, Cas7-2x.1, and Cas7-Cas5-Cas11, respectively. The RNA was labelled at the 5’ end with an IRD800 label.

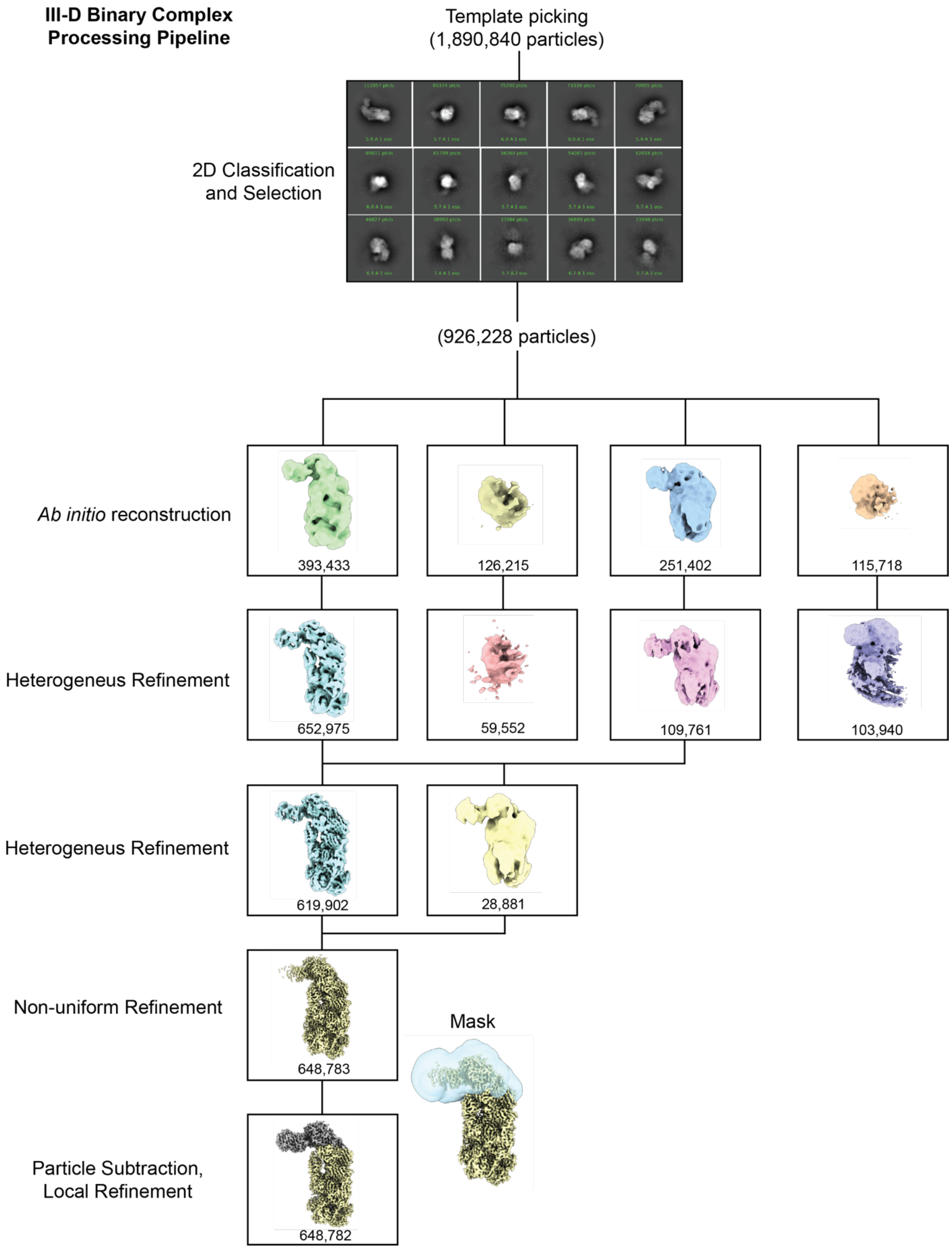


**Supplementary Figure 14: Cryo-EM data processing workflow for the type III-Dv binary complex.**

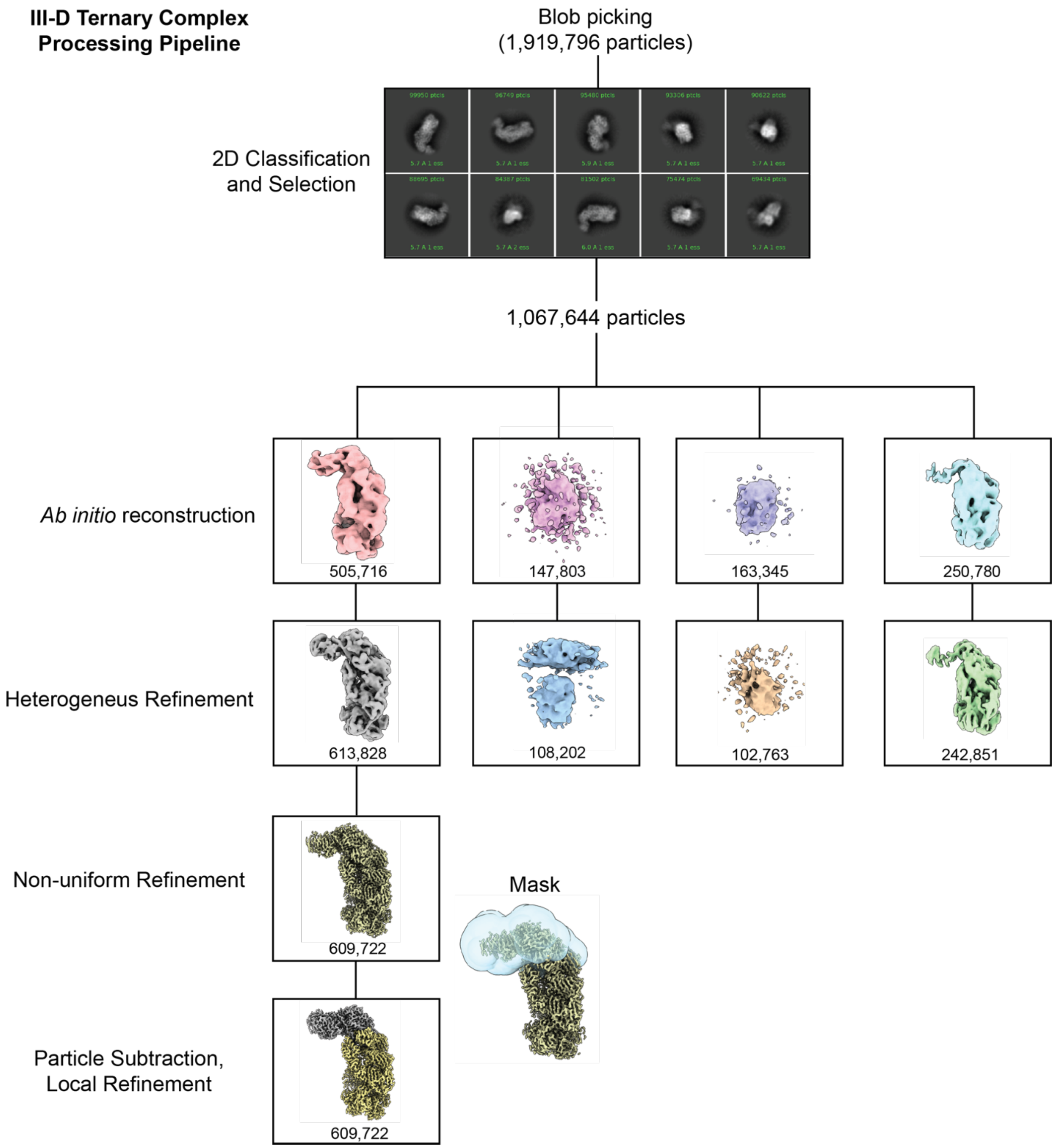


**Supplementary Figure 15: Cryo-EM data processing workflow for the type III-Dv ternary complex.**
